# H2A ubiquitination is essential for Polycomb Repressive Complex 1-mediated gene regulation in *Marchantia polymorpha*

**DOI:** 10.1101/2021.04.27.441584

**Authors:** Shujing Liu, Minerva S. Trejo-Arellano, Yichun Qiu, D. Magnus Eklund, Claudia Köhler, Lars Hennig

**Affiliations:** Department of Plant Biology, Swedish University of Agricultural Sciences and Linnean Center for Plant Biology, Uppsala 75007, Sweden; Department of Cell and Developmental Biology, John Innes Centre, Norwich NR4 7UH, UK; Department of Plant Ecology and Evolution, Evolutionary Biology Centre, Uppsala University, Uppsala 75236, Sweden

## Abstract

Polycomb repressive complex 1 (PRC1) and PRC2 are chromatin regulators maintaining transcriptional repression. The deposition of H3 lysine 27 tri-methylation (H3K27me3) by PRC2 is known to be required for transcriptional repression, whereas the contribution of H2A ubiquitination (H2Aub) in the Polycomb repressive system remains unclear in plants. We directly tested the requirement of H2Aub for gene regulation in *Marchantia polymorpha* by generating point mutations in H2A that prevent ubiquitination by PRC1. These mutants show reduced H3K27me3 levels on the same target sites as mutants defective in PRC1 subunits MpBMI1 and the homolog MpBMI1L, revealing that PRC1-catalyzed H2Aub is essential for Polycomb system function. Furthermore, by comparing transcriptome data between mutants in *MpH2A* and *MpBMI1/1L*, we demonstrate that H2Aub contributes to the PRC1-mediated transcriptional level of genes and transposable elements. Together, our data demonstrate that H2Aub plays a direct role in H3K27me3 deposition and is required for PRC1-mediated transcriptional changes in both genic and intergenic regions in *Marchantia*.

## Background

Polycomb group (PcG) proteins are evolutionarily conserved epigenetic regulators which maintain transcriptional gene repression in essential cellular and developmental processes in eukaryotes [1–4]. PcG proteins typically belong to one of the two functionally distinct multi-protein complexes: Polycomb Repressive Complex 1 (PRC1) and PRC2. PRC1 promotes chromatin compaction and catalyzes mono-ubiquitination on histone 2A (H2Aub) mainly at lysine 119 in mammals, lysine 118 in *Drosophila*, and lysine 121 in *Arabidopsis* [4–8], whereas PRC2 tri-methylates histone 3 at lysine 27 (H3K27me3) [9–12]. The catalytic core of the mammalian PRC1 is composed of the E3 ubiquitin ligases RING1A or RING1B and one of six Polycomb RING finger (PCGF) proteins [13–15], while in *Drosophila* it consists of RING1 (encoded by the *Sce* gene) and one of two PCGF proteins: Psc or Su(z)2 [16–18]. The *Arabidopsis* PRC1 core includes AtRING1A or AtRING1B and one of the three AtBMI1s (homologs of PCGF4) [6,19–21].

Previous studies on the Polycomb repressive system in *Drosophila* and mammals first proposed a PRC2-initiated hierarchical model where PRC2 establishes H3K27me3, which is then recognized by chromodomain-containing subunits of the canonical PRC1 (cPRC1). Nevertheless, later studies found this classical hierarchical model not sufficient to explain the Polycomb repressive system [13,22,23]. Instead, it was found that non-canonical PRC1 (ncPRC1) lacking chromodomain-containing subunits can recruit PRC2 and establish stable Polycomb repressive domains [24–27]. This data point that the PRC1 catalytic function is required for PRC2 recruitment, which was supported by recent work showing that in mouse embryonic stem cells (ESCs), loss of RING1B catalytic activity largely phenocopies the complete removal of the RING1B protein [28,29]. Nevertheless, whether this is a generally applicable concept remains to be established. In *Drosophila*, H2AK118ub seems not required for repression of Polycomb target genes during the early stages of embryo development and PRC2 binding to chromatin requires PRC1 but not H2Aub [30,31]. Similarly, during neuronal fate restriction in mouse, PRC1 repression was shown to function independently of ubiquitination of [32]. This data suggest that there are developmental context-specific differences in the functional requirement of the catalytic activity of PRC1.

PRC1-catalyzed H2Aub has been intensively studied in *Arabidopsis thaliana*. H2Aub level, H3K27me3 incorporation and chromatin accessibility were shown to be affected by the depletion of components of PRC1 [33–35]. Nevertheless, it remains unclear thus far whether H2Aub is required for H3K27me3 targeting. PRC1 is composed of multiple proteins that engage in interactions with PRC2 components. Thus, AtRING and AtBMI1 in PRC1 can interact with LHP1, which co-purifies with PRC2 [19,20,36], suggesting that PRC1 rather than H2Aub promotes H3K27me3 by interacting and recruiting PRC2 to chromatin. H2Aub is associated with permissively accessible chromatin and the average transcription levels of only-H2Aub marked genes are higher than that of H2Aub/H3K27me3 and only-H3K27me3 marked genes in *Arabidopsis* [33,35]. Consistently, removal of H2Aub is required for stable repression of Polycomb target genes [34]. Together, based on current studies, the direct role of H2Aub in transcriptional regulation remains unclear. To explore the functional role of H2Aub, we generated H2Aub mutants by replacing the endogenous H2A by H2A variants with mutated lysines in the liverwort *Marchantia polymorpha*.

*Marchantia* shares many signaling pathways with *Arabidopsis* and other seed plants [37]. Together with its low genetic redundancy and possibilities to easily generate mutants, *Marchantia* is an ideal plant model to study the evolutionarily conserved Polycomb system. There is only a single gene encoding canonical H2A in *Marchantia*, compared to four genes in *Arabidopsis* [38]. We generated lysine to arginine substitutions in H2A on residues K115/116/119 and demonstrate that all three lysines are ubiquitinated *in vivo* and likely have redundant functions. We furthermore show that H2Aub mediates H3K27me3 incorporation in both genic and intergenic regions in *Marchantia* and reveal that H2Aub is essential for both, PRC1-mediated transcriptional activation and silencing.

## Results

### H2Aub mediates H3K27me3 deposition on Polycomb target sites in genic and intergenic regions

In mutants of PRC1 components, decreased H2Aub correlates with reduced H3K27me3 [33,35]. To elucidate the functional requirement of H2Aub to induce H3K27me3 incorporation, we generated H2Aub depleted lines by introducing point mutations in the potential ubiquitination sites of canonical MpH2A (Fig. S1a). We co-transformed the point mutated *H2A* variants and a CRISPR construct designed to knock out the endogenous *H2A* (Fig. S1b, S1c). Lysine 120 (K120) and K121 of *Arabidopsis* H2A were shown to be ubiquitinated by AtBMI1 *in vitro* [6,20]; corresponding to K115 and K116 in MpH2A (Fig. 1a). In *Drosophila*, mutations of four close lysine sites (K117, K118, K121 and K122) are required to abolish total H2Aub [30]. We therefore generated *Mph2a;H2AK115R/K116R* and *Mph2a;H2AK119R* mutants (jointly referred to as *h2a_ub* mutants) by substituting the C-terminal lysine residues K115 and K116 or K119 of MpH2A with arginine. We failed to obtain *Mph2a* mutants expressing *H2A* variants with all three point mutations, indicating that the three lysine sites of MpH2A are functionally redundant. The global H2Aub level was strongly decreased in lines of *Mph2a;H2AK115R/K116R* and *Mph2a;H2AK119R* mutants compared to wild type (WT) (Fig. 1b). Nevertheless, there was a remaining H2Aub signal in both mutant lines, indicating that all lysine residues can be ubiquitinated *in vivo* and likely act redundantly. The most obvious defects of *h2a_ub* mutants were downward curled edges of the thallus that grew into the growth media and decreased gemmae dormancy compared to WT (Fig. 1c–1h).

**Fig. 1.**
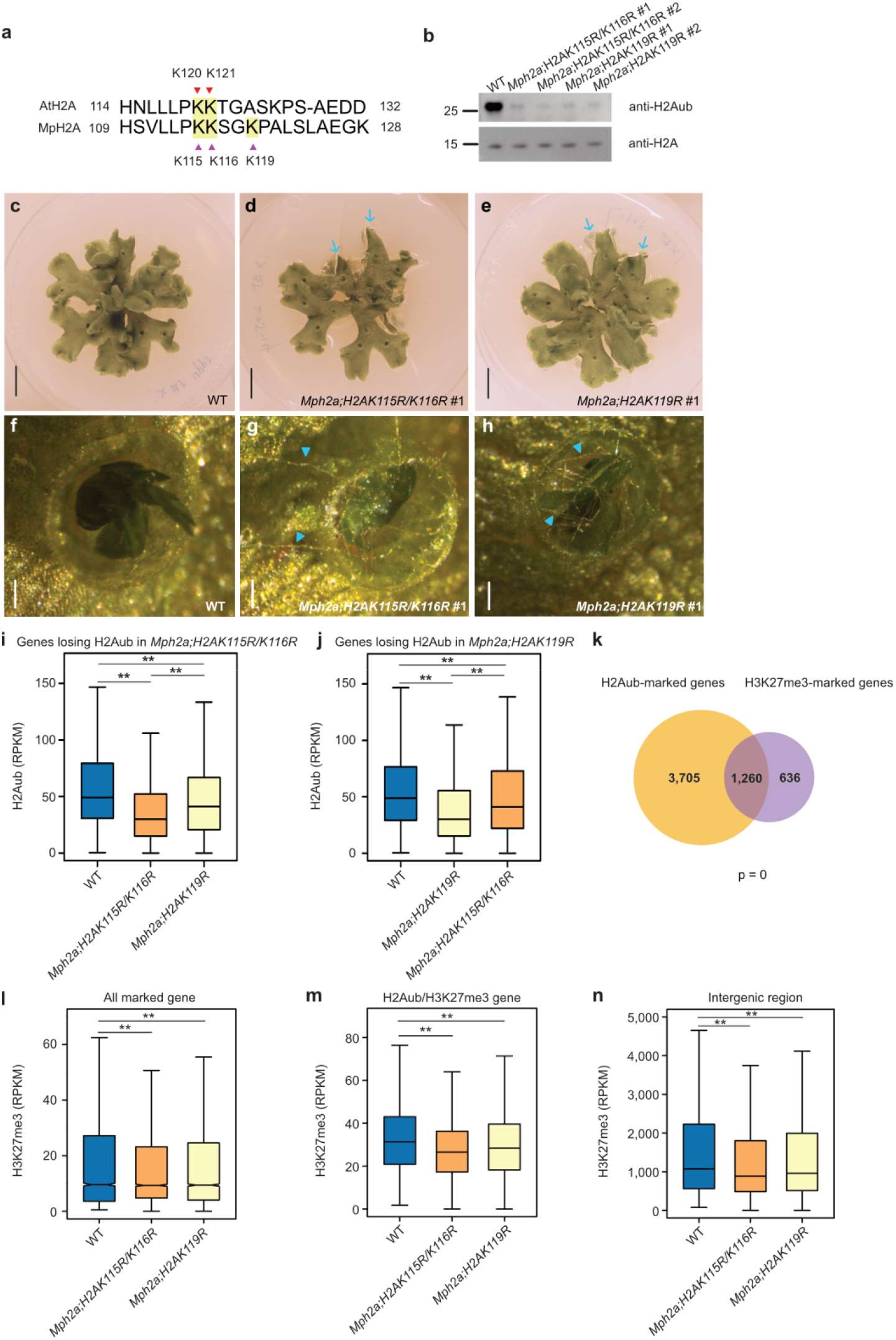
H2Aub directly contributes to the deposition of H3K27me3. a. Lysine residues in *Arabidopsis* and *Marchantia* H2A. Lysine 120 (K120) and K121 of H2A in *Arabidopsis* and potential ubiquitination sites (K115, K116 and K119) in H2A of *Marchantia* are highlighted. b. Western blot of bulk H2Aub and H2A in wild type (WT), *Mph2a;H2AK115R/K116R* #1, *Mph2a;H2AK115R/K116R* #2, *Mph2a;H2AK119R* #1 and *Mph2a;H2AK119R* #2 mutants. c-e. Phenotypes of 28 day-old WT, *Mph2a;H2AK115R/K116R* #1 and *Mph2a;H2AK119R* #1 mutants. Blue arrows point to downward curled edges of the thallus. Scale bars, 1 cm. f-h. Gemma cup phenotypes of WT, *Mph2a;H2AK115R/K116R* #1 and *Mph2a;H2AK119R* #1 mutants. Blue arrowheads point at visible rhizoids produced by non-dormant gemmae in the mutants. Scale bars, 0.05 cm. i. Boxplots showing H2Aub levels (RPKM, reads per kilobase per million mapped reads) of genes losing H2Aub in *Mph2a;H2AK115R/K116R* in WT, *Mph2a;H2AK115R/K116R* and *Mph2a;H2AK119R* mutants. j. Boxplots showing H2Aub levels of genes losing H2Aub in *Mph2a;H2AK119R* in WT, *Mph2a;H2AK119R* and *Mph2a;H2AK115R/K116R* mutants. k. Venn diagram showing overlap of H2Aub-marked genes and H3K27me3-marked genes in wild type (WT). Significance was tested using a Hypergeometric test. l, m. Boxplots showing H3K27me3 levels of all marked genes (l) and H2Aub/H3K27me3 genes (m) in WT, *Mph2a;H2AK115R/K116R* and *Mph2a;H2AK119R* mutants. H2Aub and H3K27me3 levels were calculated as the average RPKM from 1 kb upstream of the transcriptional start to the transcriptional start of genes. n. Boxplot showing H3K27me3 levels in intergenic regions marked by H2Aub and H3K27me3 in WT, *Mph2a;H2AK115R/K116R* and *Mph2a;H2AK119R* mutants. Boxes show medians and the interquartile range, and error bars show the full range excluding outliers. **, p < 0.01 (Wilcoxon test).

To understand the connection between H2Aub and H3K27me3 in the Polycomb repressive system, we generated ChIP-seq data for H3, H2Aub and H3K27me3 in WT, *Mph2a;H2AK115R/K116R* #1 and *Mph2a;H2AK119R* #1 mutants. To validate our ChIP-seq data, we compared the H3K27me3 peaks in our WT with previously published data [39]. The majority of peaks overlapped between both datasets (Fig. S2), supporting the quality of our data. We found that genes with decreased H2Aub in the *Mph2a;H2AK115R/K116R* mutant had also decreased H2Aub levels in the *Mph2a;H2AK119R* mutant, but to a lesser extent (Fig. 1i). Conversely, genes with reduced H2Aub level in the *Mph2a;H2AK119R* mutant were less affected in the *Mph2a;H2AK115R/K116R* mutant (Fig. 1j), supporting the notion that all three lysine residues of MpH2A are targeted by ubiquitination. To test whether H3K27me3 deposition is affected upon H2Aub depletion, we analyzed H3K27me3 on genes marked by either H2Aub or H3K27me3 (all marked genes, all genes in Fig. 1k) and genes marked by both H2Aub and H3K27me3 (H2Aub/H3K27me3, overlapped genes in Fig. 1k). We found H3K27me3 levels to be decreased on all marked genes and a more pronounced decrease on H2Aub/H3K27me3 genes in both *h2a_ub* mutants compared to WT (Fig. 1l, 1m), revealing that H2Aub is essential for H3K27me3 deposition.

In *Marchantia*, 60% of H3K27me3 peaks are present in intergenic regions [39]; however, the location of H2Aub peaks remains to be explored. Out of 6575 H2Aub peaks identified in WT, about 20% mapped to intergenic regions, while most of the H2Aub peaks were located in gene body or promoter regions (Fig. S3). We found intergenic regions covered by H2Aub and H3K27me3 also had decreased H3K27me3 levels in both *h2a_ub* mutants (Fig. 1n), revealing that H2Aub is required in intergenic regions to recruit H3K27me3 in plants.

### H2Aub contributes to transcriptional activation

Although many transcriptionally active genes are marked with H2Aub and H2Aub is associated with a permissive chromatin state in *Arabidopsis* [33–35], it is unknown whether H2Aub is required for gene activation. We noted that in *h2a_ub* mutants there were more downregulated than upregulated genes (Fig. 2a, 2b), suggesting that H2Aub has an activating role for gene expression in *Marchantia*. Both upregulated and downregulated genes in *h2a_ub* mutants were enriched for genes with only-H2Aub and H2Aub/H3K27me3 (Fig. 2c). We tested the H2Aub level on upregulated and downregulated genes in *h2a_ub* mutants and found that H2Aub levels were decreased in the promoter regions of both gene categories (Fig. 2d–2g), supporting that H2Aub contributes to gene repression as well as activation. We also found more downregulated than upregulated TEs in *h2a_ub* mutants (Fig. 2h, 2i). There were 559 TEs marked with only-H2Aub, 1059 with H2Aub/H3K27me3 and 5728 with only-H3K27me3 (Fig. S4). Among those, the only-H2Aub and H2Aub/H3K27me3 marked TEs had a significantly higher number of highly transcriptionally active TEs than the only-H3K27me3 marked TEs (Fig. 2j), pointing that H2Aub has the potential to activate TE expression and that removal of H2Aub is required for stable repression.

**Fig. 2.**
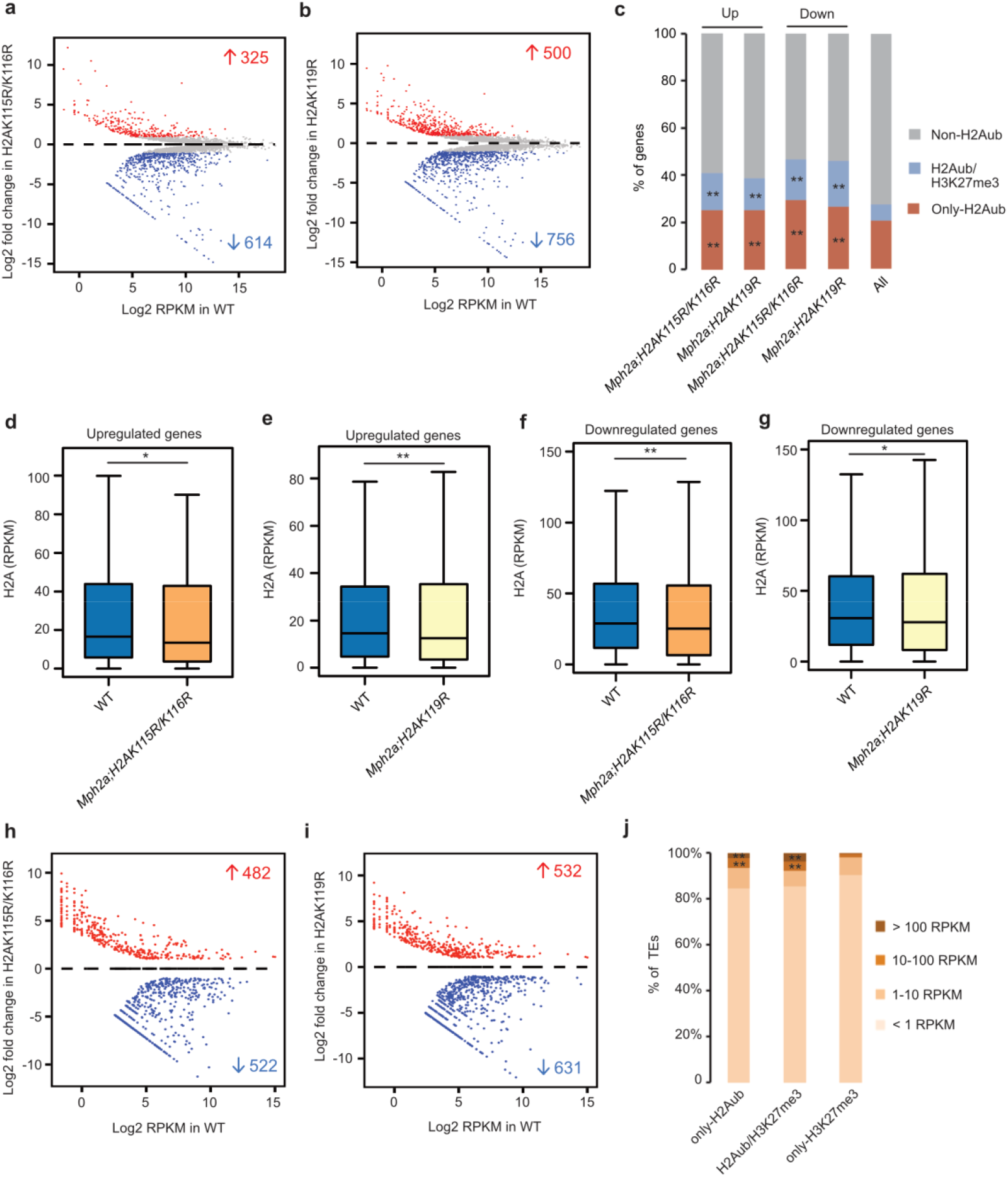
H2Aub is required for gene activation. a, b. MA plots showing differential gene expression (Log2 fold change) in *Mph2a;H2AK115R/K116R* (a) and *Mph2a;H2AK119R* (b) mutants compared to wild type (WT). Significant changes are marked in red (log2 fold change ≥ 1 and adjusted p value ≤ 0.05) and blue (log2 fold change ≤ −1 and adjusted p value ≤ 0.05). c. Presence of only-H2Aub or H2Aub/H3K27me3 marks on upregulated (Up) and downregulated (Down) genes in *Mph2a;H2AK115R/K116R* and *Mph2a;H2AK119R* mutants. Presence of modifications is based on their distribution in WT. **, p < 0.01 (Hypergeometric test). d, e. Boxplots showing the H2Aub level on upregulated genes in *Mph2a;H2AK115R/K116R* (d) and *Mph2a;H2AK119R* (e) mutants. f, g. Boxplots showing H2Aub level on downregulated genes in *Mph2a;H2AK115R/K116R* (f) and *Mph2a;H2AK119R* (g) mutants. H2Aub levels were calculated as the average RPKM from 1 kb upstream of the transcriptional start to the transcriptional start of genes. oxes show medians and the interquartile range, and error bars show the full range excluding outliers. **, p < 0.01 (Wilcoxon test). h, i. MA plots showing differential expression of transposable elements (TEs) (Log2 fold change) in *Mph2a;H2AK115R/K116R* (h) and *Mph2a;H2AK119R* (i) mutants. Significant gene expression changes are marked in red (log2 fold change ≥ 1 and adjusted p value ≤ 0.05) and blue (log2 fold change ≤ −1 and adjusted p value ≤ 0.05). j. Expression level distribution of transposable elements (TEs) marked by only-H2Aub, H2Aub/H3K27me3 and only-H3K27me3 in WT. **, p < 0.01 (Hypergeometric test).

### MpBMI1/1L regulate morphological development of *Marchantia* and are involved in gene repression and activation

In *Arabidopsis*, AtBMI1s have been shown to be involved in H2A ubiquitination [6,20,21,33]. To explore the extent to which MpBMI/1L function relies on H2Aub, we generated MpBMI1 knock out mutants by CRISPR/Cas9 in *Marchantia*. Using the AtBMI1A protein sequence as a query in a protein blast, we identified two genes (*Mp7g12670* and *Mp6g09730* in the MpTak1 v5.1 annotation) encoding AtBMI1 homologs in *Marchantia*. MpBMI1, encoded by *Mp7g12670*, contains an N-terminal RING finger domain and a C-terminal ubiquitin-like (RAWUL) domain [40], while *Mp6g09730* encodes for a protein named MpBMI1-LIKE (MpBMI1L) that only has a C-terminal RAWUL domain. The RAWUL domain is involved in protein-protein interaction and oligomerization of BMI1, which is essential for H2Aub activity of PRC1 in mammals [41–44]. We generated *Mpbmi1/1l* double knockout mutants by CRISPR/Cas9 and obtained combinations of double mutants with different mutations at the Cas9 target sites (mutant information shown in Fig. S5, S6). Combinations of strong mutant alleles for both genes *Mpbmi1-1/Mpbm1l-1* (named *Mpbmi1/1l*#1, Fig. S5a) and *Mpbmi1-2/Mpbm1l-2* (named *Mpbmi1/1l* #2, Fig. S5b) caused strongly reduced growth rates and substantial size-reduction of gemma cups that contained only few and smaller gemmae compared to WT (Fig. 3a–3c and 3f–3h). The slightly more severe size reduction in *Mpbmi1/1l* #1 compared to *Mpbmi1/1l* #2 is likely due to the different extent of deletions and insertions caused by CRISPR/Cas9 in these two lines (Fig. S5). We also obtained one double mutant with a strong *Mpbmi1-3* allele and a weak *Mpbmi1l-3* allele (named *Mpbmi1/1l* #3, Fig. S6a) and one *Mpbmi1l-4* single mutant (Fig. S6b), which showed weakly reduced growth rates compared to WT (Fig. 3d, 3e). The fact that *Mpbmi1/1l* mutants had a more severe phenotype than *h2a_ub* mutants is consistent with the proposed redundant function of ubiquitination on K115/K116 and K119 in H2A. Western blot analysis revealed a global decrease of H2Aub in *Mpbmi1/1l* #1, *Mpbmi1/1l* #2 and *Mpbmi1/1l* #3 double mutants compared to WT (Fig. 3i), with a more pronounced reduction in the *Mpbmi1/1l* #1 mutant combination. We therefore used *Mpbmi1/1l* #1 in subsequent analyses. The residual H2Aub signal in the *Mpbmi1/1l* mutants possibly reflects remaining functional activity of MpBMI1/1L generated in the mutants. Alternatively, MpRING proteins have low functional activity in the absence of MpBMI1/1L. We found 2085 genes being upregulated and 1023 genes being downregulated in the *Mpbmi1/1l* mutants (Fig. 3j), suggesting that MpBMI1/1L mainly function as repressors, but also possibly as activators. We analyzed H2Aub levels on the deregulated genes in the *Mpbmi1/1l* mutants and found that H2Aub level significantly decreased on both upregulated genes and downregulated genes (Fig. 3k), implying that H2Aub is required for MpBMI1/1L-mediated gene silencing and activation. Accordingly, H2Aub marked genes were enriched in both upregulated and downregulated genes in *Mpbmi1/1l* mutants (Fig. 3l).

**Fig. 3.**
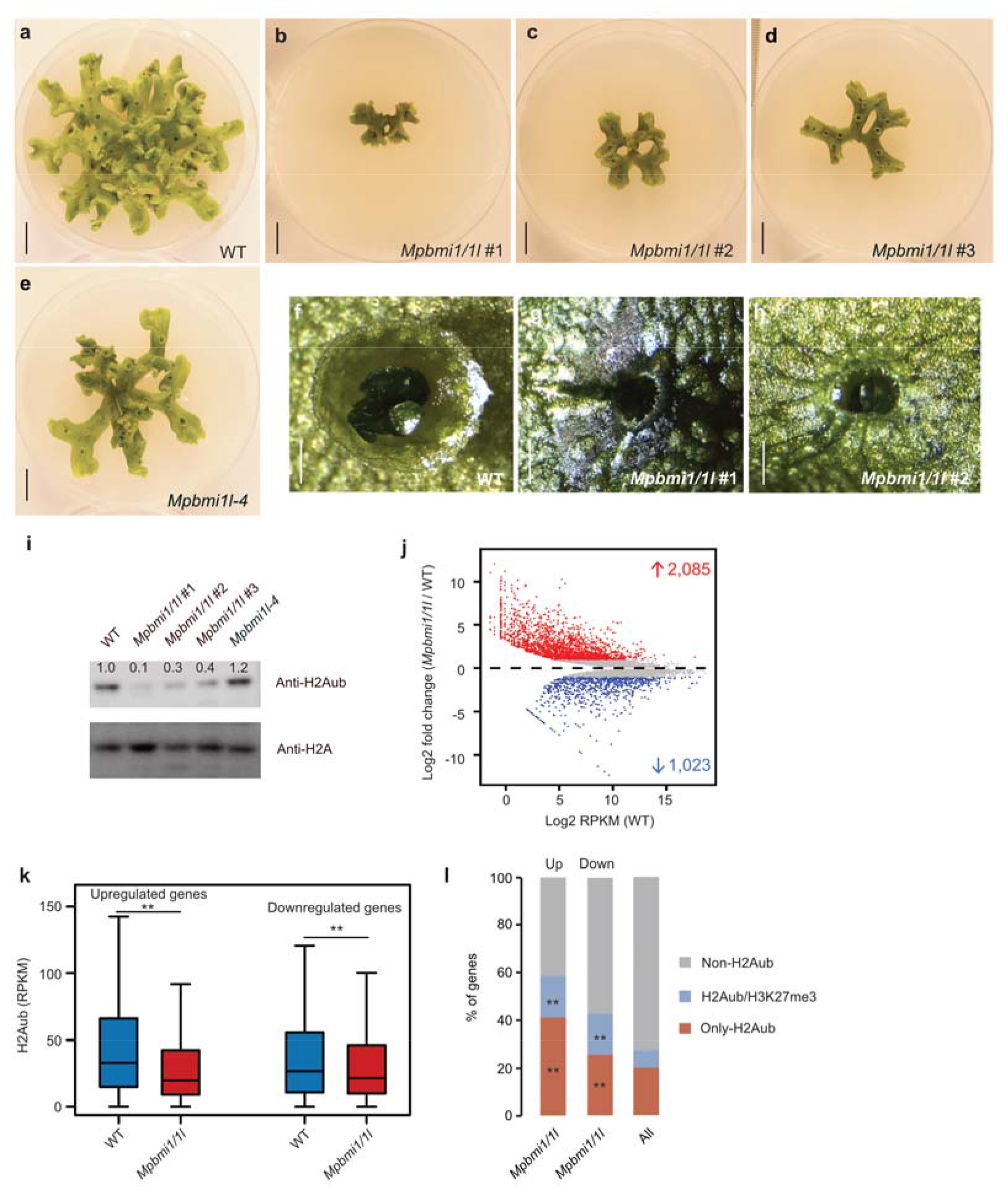
MpBMI1/1L affect morphological development of *Marchantia* and regulate gene silencing and activation. a-e. Morphological phenotypes of 35 day-old wild type (WT), *Mpbmi1/1l* #1, *Mpbmi1/1l* #2, *Mpbmi1/1l* #3 and *Mpbmi1l-4* lines. Scale bars: 1 cm. f-h. Gemma cups of 35 day-old WT, *Mpbmi1/1l* #1 and *Mpbmi1/1l* #2 lines. Scale bars: 0.1 cm. i. Western blot showing the bulk H2Aub levels in WT, *Mpbmi1/1l* #1, *Mpbmi1/1l* #2, *Mpbmi1/1l* #3 and *Mpbmi1l-4* mutants. j. MA plot showing differential gene expression (Log2 fold change) in *Mpbmi1/1l* #1 mutants compared to WT. Significant gene expression changes are marked in red (log2 fold change ≥ 1 and adjusted p value ≤ 0.05) and blue (log2 fold change ≤ −1 and adjusted p value ≤ 0.05). i. Boxplot showing the H2Aub level (RPKM, reads per kilobase per million mapped reads) of upregulated and downregulated genes in *Mpbmi1/1l* #1 mutants. H2Aub levels were calculated as the average RPKM from 1 kb upstream of the transcriptional start to the transcriptional start of genes. Boxes show medians and the interquartile range, and error bars show the full range excluding outliers. **, p < 0.01 (Wilcoxon test). j. Percent of upregulated (Up) and downregulated (Down) genes marked by only-H2Aub and H2Aub/H3K27me3 in *Mpbmi1/1l* #1 mutants. **, p < 0.01 (Hypergeometric test).

### Ubiquitination of H2AK115/K116 and H2AK119 is required for PRC1-mediated gene and transposable element expression

To test whether impaired H2A ubiquitination and loss of PRC1 function has similar consequences, we compared transcriptome data of the *Mph2a;H2AK115R/K116R* and *Mph2a;H2AK119R* mutants with that of the *Mpbmi1/1l* mutants. Both *h2a_ub* mutants shared a significant number of upregulated genes (Fig. 4a) and we also found a significant overlap of upregulated genes between the *Mpbmi1/1l* mutants and *h2a_ub* mutants (Fig. 4b, 4c) as well as between all mutants (Fig. 4d). Similarly, a significant number of downregulated genes overlapped between two *h2a_ub* mutants and *Mpbmi1/1l* mutants (Fig. 4e–4h). Genes commonly upregulated in *Mpbmi1/1l* and *Mph2a;H2AK115R/K116R* or *Mph2a;H2AK119R* mutants were more strongly upregulated in *Mpbmi1/1l* than in the *h2a_ub* mutants (Fig. 4i, 4j), supporting the idea that mono-ubiquitination on H2AK115/K116 and H2AK119 is functionally redundant and mediated by MpBMI1/1L. Commonly downregulated genes in *h2a_ub* and *Mpbmi1/1l* mutants were expressed at similar low levels in *h2a_ub* and *Mpbmi1/1l* mutants (Fig. 4k, 4l), consistent with the idea that H2Aub is required for PRC1-mediated gene activation. Furthermore, it suggests that ubiquitination of H2AK115/K116 and H2AK119 is not functionally redundant in activating gene expression. GO enrichment analyses of upregulated genes overlapping between *Mpbmi1/1l* and *Mph2a;H2AK115R/K116R* mutants (Fig. 4m) or *Mpbmi1/1l* and *Mph2a;H2AK119R* mutants (Fig. 4n) both showed that response pathways were over-represented. Among downregulated genes overlapped between *Mpbmi1/1l* and *h2a_ub* mutants we also found a significant enrichment for response pathway related GOs (Fig. 5a, 5b), which were nevertheless largely distinct from the enriched GO terms of upregulated genes.

**Fig. 4.**
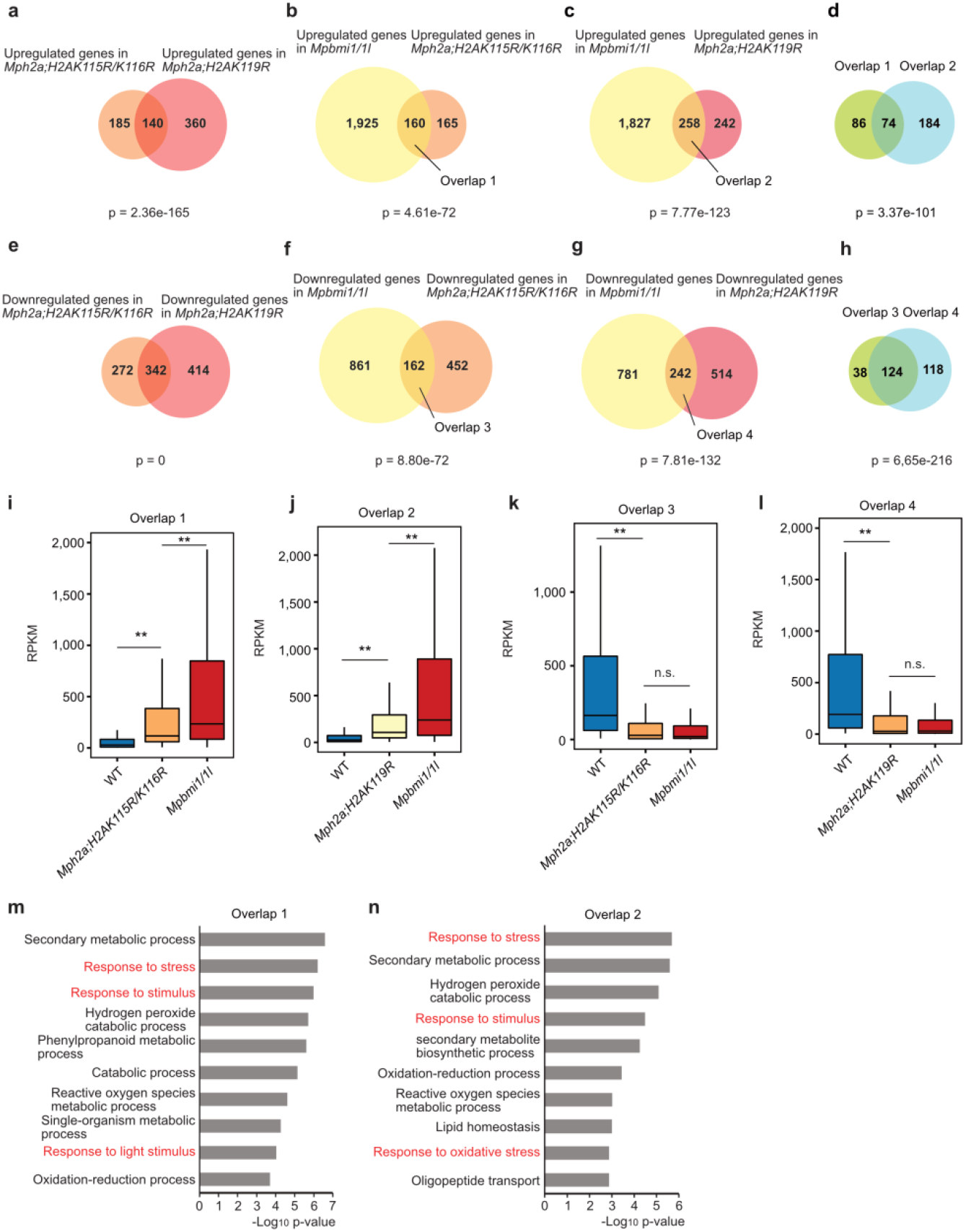
H2Aub contributes to PRC1-mediated transcriptional expression. a. Venn diagram showing overlap of upregulated genes in *Mph2a;H2AK115R/K116R* and *Mph2a;H2AK119R* mutants. Significance was tested using a Hypergeometric test. b. Venn diagram showing overlap of upregulated genes in *Mpbmi1/1l* mutants and *Mph2a;H2AK115R/K116R* mutants. c. Venn diagram showing overlap of upregulated genes in *Mpbmi1/1l* mutants and *Mph2a;H2AK119R* mutants. d. Venn diagram showing overlap of commonly upregulated genes in overlap 1 (panel b) and overlap 2 (panel c). e. Venn diagram showing overlap of downregulated genes in *Mph2a;H2AK115R/K116R* and *Mph2a;H2AK119R* mutants. f. Venn diagram showing overlap of downregulated genes in *Mpbmi1/1l* mutants and *Mph2a;H2AK115R/K116R* mutants. g. Venn diagram showing overlap of downregulated genes in *Mpbmi1/1l* mutants and *Mph2a;H2AK119R* mutants. h. Venn diagram showing overlap of commonly downregulated genes in overlap 3 (panel f) and overlap 4 (panel g). i. Boxplot showing expression level (RPKM, reads per kilobase per million mapped reads) of genes in overlap 1 (panel b) in WT, *Mph2a;H2AK115R/K116R* and *Mpbmi1/1l*. j. Boxplot showing expression level (RPKM) of genes in overlap 2 (panel c) in WT, *Mph2a;H2AK119R* and *Mpbmi1/1l*. k. Boxplot showing expression level (RPKM, reads per kilobase per million mapped reads) of genes in overlap 3 (panel f) in WT, *Mph2a;H2AK115R/K116R* and *Mpbmi1/1l*. l. Boxplot showing expression level (RPKM) of genes in overlap 4 (panel g) in WT, *Mph2a;H2AK119R* and *Mpbmi1/1l*. Boxes show medians and the interquartile range, and error bars show the full range excluding outliers. **, p < 0.01 (Wilcoxon test). m, n. Enriched GO terms of commonly upregulated genes in overlap 1 (panel b) and overlap 2 (panel c). Response pathways are marked in red.

**Fig. 5.**
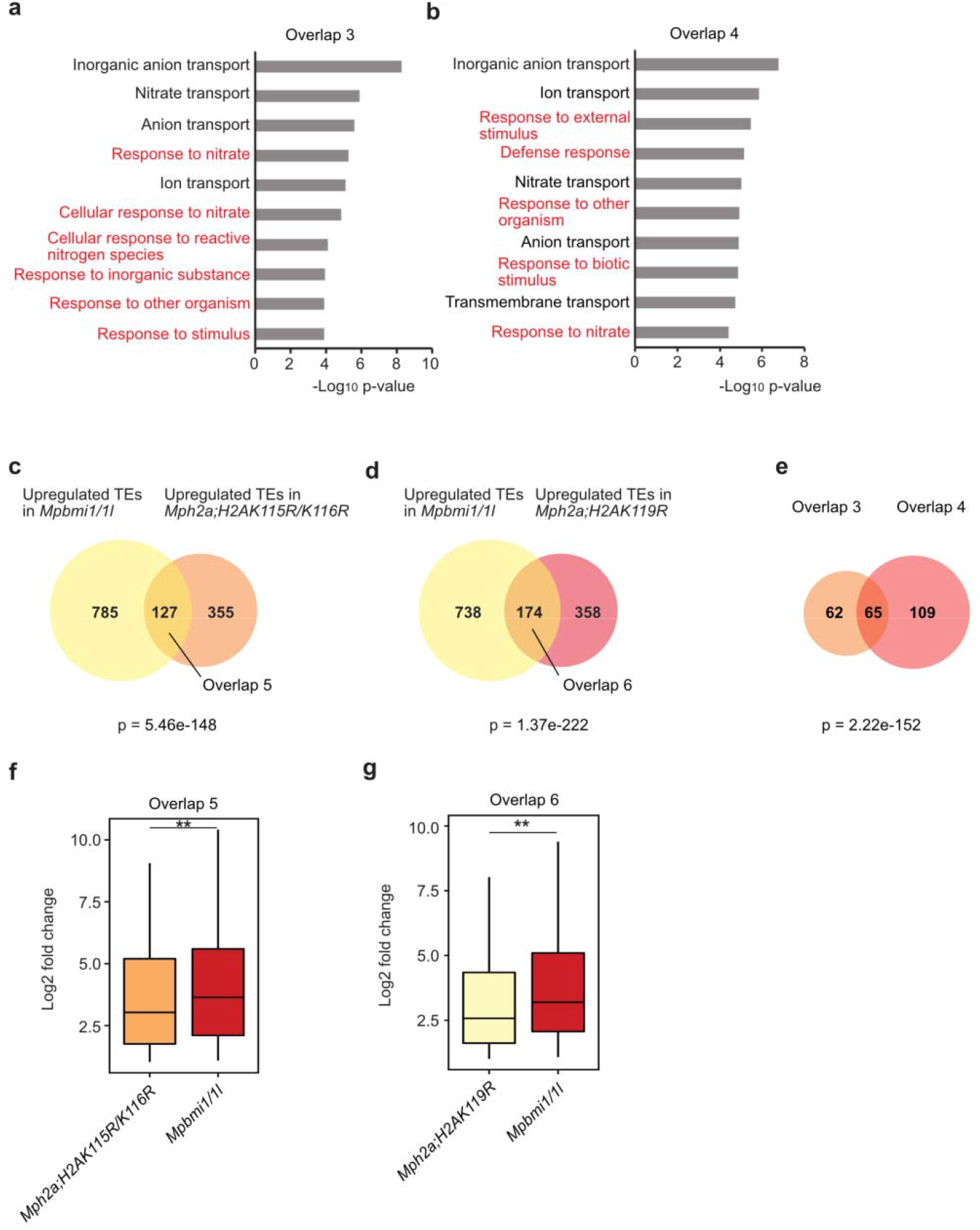
H2Aub is essential in PRC1-mediated transposable element silencing. a, b. Enriched GO terms of commonly downregulated genes in overlap 3 (panel f in Fig. 4) and overlap 4 (panel g in Fig. 4). Response pathways are marked in red. c. Venn diagram showing overlap of upregulated transposable elements (TEs) in *Mpbmi1/1l* and *Mph2a;H2AK115R/K116R* mutants. Significance was tested using a Hypergeometric test. d. Venn diagram showing overlap of upregulated TEs in *Mpbmi1/1l* and *Mph2a;H2AK119R* mutants. Significance was tested using a Hypergeometric test. e. Venn diagram showing overlap of commonly upregulated TEs in overlap 5 (panel c) and overlap 6 (panel d). f. Boxplot showing differential expression level (Log_2_ fold change) of TEs in overlap 5 (panel c) in *Mph2a;H2AK115R/K116R* and *Mpbmi1/1l* mutants compared to WT. n. Boxplot showing differential expression level (Log_2_ fold change) of TEs in overlap 6 (panel d) in *Mph2a;H2AK119R* and *Mpbmi1/1l* mutants compared to WT. Boxes show medians and the interquartile range, and error bars show the full range excluding outliers. **, p < 0.01 (Wilcoxon test).

The localization of H2Aub in intergenic regions indicated a role of this modification for transposable element (TE) repression. Indeed, we identified more than 900 upregulated TEs in *Mpbmi1/1l* mutants, of which a significant number overlapped with upregulated TEs in *h2a_ub* mutants (Fig. 5c–5e), revealing that H2Aub is critical for PRC1-mediated TE silencing. Like for genes, commonly upregulated TEs between *Mpbmi1/1l* mutants and *h2a_ub* mutants were more strongly upregulated in *Mpbmi1/1l* mutants compared to *h2a_ub* mutants (Fig. 5f, 5g), pointing at redundant regulation of TEs by mono-ubiquitination of H2AK115/K116 and H2AK119.

### H2Aub and H3K27me3 are affected in genic and intergenic regions by the depletion of MpBMI1/1L

Consistent with the effect caused by AtBMI1 depletion in *Arabidopsis* [33], the H2Aub level was significantly decreased on all marked genes as well as H2Aub/H3K27me3 genes in the *Mpbmi1/1l* mutants compared to WT (Fig. 6a, 6b and S7a). Also the H3K27me3 level was significantly decreased on all marked genes and H2Aub/H3K27me3 genes, yet not on only H3K27me3 genes in the *Mpbmi1/1l* mutants compared to WT (Fig. 6c, 6d and S7b), implying that PRC1 activity mediates H3K27me3 deposition in *Marchantia*. Consistently, genes losing H2Aub showed significantly reduced H3K27me3 levels in the *Mpbmi1/1l* mutants compared to WT (Fig. 6e). We tested whether genes losing H2Aub in the *Mpbmi1/1l* mutants belong to specific pathways. Among the top twenty significantly enriched GO terms, nine GO terms correspond to multiple response pathways to intrinsic and extrinsic stimuli (Fig. 6f), which occurred in the commonly upregulated and downregulated genes between *Mpbmi1/1l* mutants and *h2a_ub* mutants. We tested whether the connection between H2Aub and H3K27me3 was restricted to genic regions or was also present in intergenic regions. Loss of MpBMI1/1L caused a significant decrease of both, H2Aub and H3K27me3 levels in intergenic regions (Fig. 6g–6i), revealing that PRC1-mediated recruitment of PRC2 is not restricted to genic regions.

**Fig. 6.**
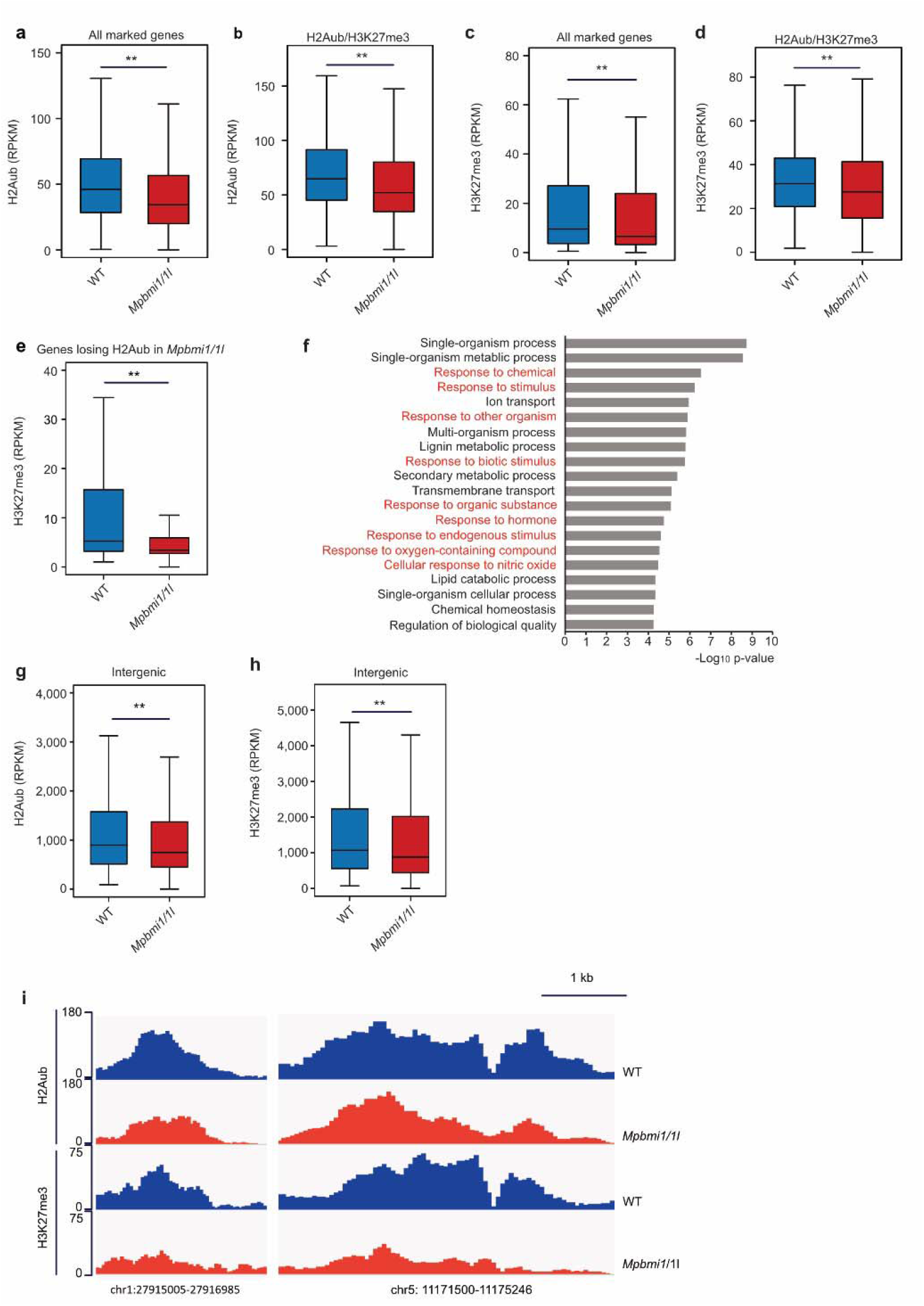
H3K27me3 is decreased on Polycomb target genes in genic and intergenic region by depletion of MpBMI1/1L. a, b. Boxplots showing H2Aub level (RPKM, reads per kilobase per million mapped reads) on all marked genes (a) and H2Aub/H3K27me3 genes (b) in WT and *Mpbmi1/1l* mutants. c, d. Boxplots showing H3K27me3 level on all marked genes (c) and H2Aub/H3K27me3 genes (d) in WT and *Mpbmi1/1l* mutants. H2Aub and H3K27me3 levels were calculated as the average RPKM from 1 kb upstream of the transcriptional start to the transcriptional start of genes. e. Boxplot showing H3K27me3 level of genes with reduced H2Aub level in *Mpbmi1/1l* mutants. f. GO terms of genes with reduced H2Aub level in *Mpbmi1/1l* mutants. Response pathways are marked in red. g, h. Boxplots showing H2Aub (g) and H3K27me3 level (h) on H2Aub/H3K27me3-occupying intergenic regions in wild type (WT) and *Mpbmi1/1l* mutants. Boxes show medians and the interquartile range, and error bars show the full range excluding outliers. **, p < 0.01 (Wilcoxon test). i. Genome browser views of two selected intergenic loci showing decreased H2Aub and H3K27me3 marks in *Mpbmi1/1l* mutants compared to WT.

## Discussion

Understanding the extent to which the function of histone modifying enzymes requires their catalytic activity is an ongoing challenge in the chromatin field. While recent work revealed that the catalytic activity of PRC1 is required in mouse ESCs [28,29], whether this requirement if evolutionary conserved, remains to be demonstrated. We found that PRC1-catalyzed H2Aub contributes to the Polycomb-mediated transcriptional repression in *Marchantia*, similar to the reported requirements in mouse ESCs [28,29]. Our study thus supports an evolutionarily conserved Polycomb mechanism in plants and animals.

Loss of MpBMI1/1L activity in *Marchantia* impaired the genome-wide deposition of H3K27me3, similar to reported findings in *Arabidopsis* [33]. Nevertheless, it was previously unknown whether the reduction of H3K27me3 in *Atbmi1* is a consequence of decreased PRC1 catalytic activity or PRC1 non-catalytic activity, since PRC1 and PRC2 were shown to interact [19,20,36]. By comparing the H2Aub deficient mutants *Mph2a;H2AK115R/K116R* and *Mph2a;H2AK119R* with *Mpbmi1/1l* mutants, we discovered that reduction of H3K27me3 levels on Polycomb target genes in *Mpbmi1/1l* mutants also occurred in *h2a_ub* mutants that evade ubiquitination, demonstrating that H2Aub directly affects H3K27me3 deposition. Previous work showed that PRC1 initiates silencing, followed by PRC2-mediated H3K27me3 that maintains stable repression in *Arabidopsis* [6,21,33,34]. Our data add support to this model and extend it by showing that the PRC1-mediated H2Aub is required for the initial PRC2-mediated repression.

In *Marchantia*, H3K27me3 is located in heterochromatic regions and marks TEs and repeats [39], contrasting its mainly genic localization in *Arabidopsis* [33]. We show that PRC1-catalyzed H2Aub is required for TE repression in *Marchantia*, revealing an ancestral role of the Polycomb system in TE repression. Similarly, in *Drosophila*, H2AK118ub is widely distributed in intergenic regions and depletion of PRC1 or PRC2 causes a genome-wide increase of transcriptional activity in intergenic regions [45].

The failure to obtain *Mph2a;H2AK115R/K116R/K119R* mutants with complete loss of H2Aub strongly suggests that H2Aub has essential functions in *Marchantia*. Similarly, H2Aub deficient *Drosophila* embryos arrest at the end of embryogenesis, indicating that the requirement of H2Aub to regulate essential biological functions is evolutionary conserved [30]. As for H2Aub deficiency, also loss of the RING1 encoding gene *Sce* causes arrest of embryo development in *Drosophila* [30]. In contrast, we found that mutants in *MpBMI1/1L* are viable and similarly, also mutants in *Arabidopsis* BMI encoding genes are viable [21]. Nevertheless, it is possible that in both systems BMI function is not completely depleted, since the *Atbmi1b* and *Atbmi1c* mutant alleles are probably not complete null alleles [6,20] and we found remaining H2Aub present in *Mpbmi1/l* mutants (Fig. 1g), pointing that the alleles have residual activity. We failed to obtain CRISPR/Cas9 mutants using guide RNAs targeting an N-terminal region in the MpBMI1, suggesting that complete loss of PRC1 function is lethal. Nevertheless, it is also possible that RING proteins can have catalytic activity independent of BMI proteins, as suggested based on *in vitro* catalytic activity of AtRING1A and AtRING1B proteins (Bratzel et al., 2010). Previous work revealed that the H2A variant H2A.Z can be ubiquitinated in *Arabidopsis* and incorporation of this modification is required for H2A.Z-mediated transcriptional repression [46]. It is possible that MpBMI1/1L also affects ubiquitination of the H2A variant H2A.Z; which could provide an alternative explanation for the more severe phenotype of *Mpbmi1/1l* mutants compared to *h2a_ub*. Due to the lack of a suitable antibody, this possibility could not be tested.

Previous work revealed that PRC1-mediated H2Aub1 is associated with chromatin responsiveness in *Arabidopsis* and that responsive genes require H2Aub to initiate PRC2 mediated repression [34,35]. At the same time, for stable gene repression H2Aub needs to be removed by the H2A deubiquitinases UBP12 and UBP13, likely because the occurrence of H2Aub allows recruitment of the H3K27me3 demethylase REF6 (Kralemann et al., 2020). The association of H2Aub with gene activation is also supported by our study, where we found more downregulated genes than upregulated genes in the *h2a_ub* mutants and a high number of downregulated genes in *Mpbmi1/l* mutants. A significant number of downregulated genes overlapped between the mutants, suggesting that the catalytic activity of PRC1 is required for gene activation. How PRC1 activity connects to gene activation remains unclear, despite several studies reporting the co-occurrence of ncPRC1 and active genes in mammalian systems [47–50]. It was proposed that the PRC1-catalytic activity may be dispensable for PRC1 function in promoting the expression of active genes in mammalian systems [51]; however, our data rather suggest that PRC1-catalyzed H2Aub is required for gene activation. We speculate that the role of H2Aub in gene activation is connected to its proposed role in recruiting REF6 [34], an exciting hypothesis that remains to be tested.

### Conclusions

In summary, we show that the ubiquitinated lysines in MpH2A act redundantly and H2Aub directly contributes to the deposition of H3K27me3 in *Marchantia*, demonstrating the determinant role of PRC1 catalysis in the Polycomb repressive system. Together with previous findings in *Arabidopsis* and mouse ESCs [28,29,33], our study supports an evolutionarily conserved Polycomb mechanism in divergent land plants and animals. Our finding strongly support a model in which the catalytic activity of PRC1 is required for PRC2-mediated gene repression and at the same time required for PRC2-independent gene activation.

## Methods

### Plant material and growth conditions

*Marchantia polymorpha* ssp. ruderalis Uppsala accession (Upp) was used as WT and for transformation [52]. Plants were grown on vented petri dishes containing Gamborg’s B5 medium solidified with 1.4% plant agar, pH 5.5, under 16/8 h photoperiod at 22°C with a light intensity of 60-70 umol m^-2^ s-^1^. Plate lids were taped to prevent loss of water.

### Generation of DNA constructs

Vectors pMpGE_En03, pMpGE010, pMpGWB401 and pMpGWB403 used in this study were previously described [53,54]. DNA fragments used to generate the guide RNAs against *MpBMI1* and *MpBMI1L* were prepared by annealing two pairs of primers (LH4513/LH4514, LH4517/LH4518, specified in table S1). The fragments were inserted into the BsaI site of pMpGE_En03 to yield pMpGE_En03-MpBMI1gRNA02 and pMpGE_En03-MpBMI1LgRNA04, respectively, and then transferred into pMpGE010 and pMpGWB401 using the Gateway LR reaction (Thermo Fisher Scientific) to generate pMpGE010_MpBMI1gRNA and pMpGWB401_MpBMI1LgRNA. Similarly, the two pairs of primers (LH4012/LH4013, LH4303/LH4304) were used to generate pMpGE_En03-MpH2AgRNA2 and pMpGE_En03-MpH2AgRNA3, which were subsequently transferred into pMpGE010 to generate pMpGE010-MpH2AgRNA2 and pMpGE010-MpH2AgRNA3, respectively. H2AK115R/K116R and H2AK119R were amplified by two pairs of primers (LH3848/LH4309 and LH3848/LH3849) and sub-cloned into the pENTR-TOPO vector (Thermo Fisher Scientific) to generate pENTR-H2AK115R/K116R and pENTR-H2AK119R, respectively. The pENTR vectors were transferred into pMpGWB403 by Gateway LR reaction to yield pMpGWB403-H2AK115R/K115R and pMpGWB403-H2AK119R. Primers used are listed in supplemental table S1.

### Generation of transgenic Marchantia polymorpha

The constructs were transformed into spores of *Marchantia* by Agrobacterium GV3101 as described previously [55]. Spores were grown in liquid Gamborg’s B5 medium with 2% sucrose, 0.1% Casamino acids and 0.03% L-Glutamine for 10 days under constant light. Agrobacteria containing constructs were grown in liquid LB with antibiotic for two days and then pelleted. The pellet was resuspended in the spore growth media with 100 mM acetosyringone and grown for 4 h at 28°C with spinning. Agrobacteria suspension was added to spores together with acetosyringone to a final concentration of 100 mM and the mixture was grown for another two days. Sporelings were plated on selection media with 200 mg/ml Timentin. Several independent primary transformants (T1 generation) were analyzed for the presence of the transgene by genomic PCR.

### Antibodies

The antibodies used were anti-H3 (07-690, Merck Millipore, Burlington, MA, USA), anti-H2Aub (#8240S, Cell signalling technology, Danvers, MA) and anti-H3K27me3 (07-449, Merck Millipore).

### Histone extraction and western blotting

Histone extraction and western blotting of 15d seedlings were performed as previously described [36].

### RNA sequencing

50 mg of 15 day-old thalli of WT, *Mpbmi1/1l*, *Mph2a;H2AK115R/K116R* and *Mph2a;H2AK119R* mutants were used for RNA extraction. RNA was extracted using the MagMAX^™^ Plant RNA Isolation Kit (Thermo Fisher Scientific) in biological triplicates. Libraries were generated using DNA-free RNA with the NEBNext^®^ Ultra^™^ II RNA Library Prep Kit for Illumina according to manufacturer’s instructions. Sequencing was performed on an Illumina HiSeq2000 in 150-bp pair-end mode at Novogene (Hong Kong).

### Transcriptome data analysis

Untrimmed reads were mapped to the *Marchantia polymorpha* MpTak1 v5.1 reference genome [39] using STAR (v2.5.3.a, [56]). Read numbers of mapping statistics are reported in supplementary table S2. Expression counts were generated using the R function summarizeOverlaps from the package HTSeq in union mode on exons from the reference transcriptome MpTak1v5.1_r1. A comparison of RPKM in RNA-seq triplicates showed high reproducibility of data in Fig. S8. Differential expression analyses were performed using the R package DESeq2 (v1.20.0, [57]). Genes with an absolute log2 fold change ≥ 1 and FDR ≤ 0.05 were considered as differentially expressed.

### H3, H2Aub1 and H3K27me3 ChIP-seq

For H3, H2Aub, and H3K27me3 ChIP-seq, WT, *Mpbmi1/1l*, *Mph2a;H2AK115R/K116R* and *Mph2a;H2AK119R* plants were grown for 15 days on B5 medium, and then about 300 mg thalli were harvested. ChIP was performed as described before [58]. In short, vacuum infiltration with formaldehyde was performed for 2×10 minutes. Crosslinking was quenched by adding glycine to a final concentration of 0.125 M under another 5-minute vacuum infiltration. Sonication of the chromatin was done for eight 30-s ON, 30-s OFF cycles. Overnight antibody binding was performed directly after sonication, followed by adding washed protein A dynabeads (Thermo Fisher Scientific) to each ChIP aliquot. De-crosslinking and subsequent DNA recovery steps were performed using the Ipure kit v2 (Diagenode, Liège, Belgium). The Ovation Ultralow Sytem V2 (NuGEN, Redwood city, CA, USA) was used for the ChIP-seq library preparation, and 150 bp paired-end sequencing was performed on the HiseqX platform at Novogene (Hong Kong). The ChIP-seq experiments were done using two biological replicates per IP, per genotype.

### Quality control and read mapping for ChIP-seq

FastQC (https://www.bioinformatics.babraham.ac.uk/projects/fastqc/) was used to examine read quality of each sample. Low quality ends (phred of <20) and adapter sequences were removed with Trimmomatic (v0.39, [59]). Reads with low average quality were also discarded (phred < 28). For all experiments, reads were mapped to the *M. polymorpha* reference genome MpTak1v5.1 using bowtie2 (v2.3.5.1; [60]). Details on read numbers can be found in supplemental table S3. Genome sequence and gene annotation data were downloaded from the *Marchantia* website (marchantia.info).

### Peak calling

H2Aub and H3K27me3 aligned .sam files were imported into Homer [61]. Duplicated mappings were removed using Homer. Peak calling was done using Homer with histone style settings and using histone H3 as the background to control for nucleosome occupancy. The same analysis was performed with the published WT H3K27me3 data (SRA: PRJNA553138) in *Marchantia* [39]. The peak tag counts were generated by Homer. A comparison of normalized peak tag counts (RPKM) by Homer in ChIP-seq replicates showed high reproducibility of data in Fig. S9. Only peaks present in both replicates of ChIP-seq data were considered as real peaks and retained for subsequent analyses. Peaks were correlated with a gene when the peaks were located at any region of this gene or at most 2 kb upstream of its transcription start site. Lists of genes defined by the presence of H2Aub and H3K27me3 are shown in supplemental table S4. Intergenic regions covered by H2Aub and H3K27me3 are listed in supplemental table S5. The statistical comparison of differential peak tag counts was performed with DEseq2 package in R using the raw tag counts outputs from Homer. Peaks with the adjusted p-value (FDR) <□0.05 were considered as differentially changed peaks.

### Peak visualization

Peak profiles were visualized by the Integrative Genome Viewer (IGV) [62]. Bigwig files were outputted from “bamCoverage” function in deepTools [63] using Reads Per Kilobase Million (RPKM) as normalization parameter. The Bigwig files were further used in the “computMatrix” function in deepTools with the “scale-regions” as setting parameter to generate H2Aub and H3K27me3 matrix on genes from 3 kb upstream of the transcriptional start to 3 kb downstream of the transcriptional end of genes, with a bin size of 50 bp. H2Aub and H3K27me3 scores (RPKM) of genes in boxplots were calculated as the average RPKM from 2 kb upstream of the transcriptional start to the transcriptional end of genes. H2Aub and H3K27me3 levels (RPKM) in intergenic region in boxplots were normalized tag counts about intergenic peaks generated from reads files (.sam file) during peak calling by Homer.

### GO analysis

GO analyses were performed on the *Arabidopsis* homologs of *Marchantia* genes. The homologs of *Marchantia* genes in *Arabidopsis* were retrieved from PLAZA 4.0 DICOT, inferred by the Best-Hits-and-Inparalogs (BHIF) approach ([64], https://bioinformatics.psb.ugent.be/plaza/versions/plaza_v4_dicots/, supplemental table S6). GO term enrichment was performed in PLAZA 4.0 DICOT.

## Acknowledgements

The computations of the RNA-seq data were performed on resources provided by SNIC through the Uppsala Multidisciplinary Center for Advanced Computational Science (UPPMAX) under Project SNIC 2017/7-6. We would like to thank Frédéric Berger (GMI) for kindly providing *Marchantia* H2A antibodies.

## Author contributions

L.H., C.K. and S.L. conceived the study. S.L. performed the experimental work. S.L., M.S.T.-A. and Y.Q. performed the computational analysis. D.M.E. provided support to the experimental work. S.L. and C.K. interpreted the data and wrote the manuscript. All authors approved the final version of the manuscript.

## Availability of data and materials

The datasets supporting the conclusions of this article are available in the Gene Expression Omnibus (GEO) with the accession number GSE164394.

## References

1. Di Croce L, Helin K. Transcriptional regulation by Polycomb group proteins. Nat Struct Mol Biol. Nature Publishing Group; 2013;20:1147–55.

2. Schwartz YB, Pirrotta V. A new world of Polycombs: unexpected partnerships and emerging functions. Nat Rev Genet. Nature Publishing Group; 2013;14:853–64.

3. Mozgova I, Hennig L. The Polycomb Group Protein Regulatory Network. Annu Rev Plant Biol. Annual Reviews; 2015;66:269–96.

4. Schuettengruber B, Bourbon H-M, Di Croce L, Cavalli G. Genome Regulation by Polycomb and Trithorax: 70 Years and Counting. Cell. 2017;171:34–57.

5. Francis NJ, Kingston RE, Woodcock CL. Chromatin Compaction by a Polycomb Group Protein Complex. Science. American Association for the Advancement of Science; 2004;306:1574–7.

6. Bratzel F, López-Torrejón G, Koch M, Del Pozo JC, Calonje M. Keeping Cell Identity in Arabidopsis Requires PRC1 RING-Finger Homologs that Catalyze H2A Monoubiquitination. Curr Biol. 2010;20:1853–9.

7. Eskeland R, Leeb M, Grimes GR, Kress C, Boyle S, Sproul D, et al. Ring1B Compacts Chromatin Structure and Represses Gene Expression Independent of Histone Ubiquitination. Mol Cell. 2010;38:452–64.

8. Xiao J, Wagner D. Polycomb repression in the regulation of growth and development in Arabidopsis. Curr Opin Plant Biol. 2015;23:15–24.

9. Cao R, Wang L, Wang H, Xia L, Erdjument-Bromage H, Tempst P, et al. Role of Histone H3 Lysine 27 Methylation in Polycomb-Group Silencing. Science. American Association for the Advancement of Science; 2002;298:1039–43.

10. Czermin B, Melfi R, McCabe D, Seitz V, Imhof A, Pirrotta V. Drosophila Enhancer of Zeste/ESC Complexes Have a Histone H3 Methyltransferase Activity that Marks Chromosomal Polycomb Sites. Cell. 2002;111:185–96.

11. Kuzmichev A, Nishioka K, Erdjument-Bromage H, Tempst P, Reinberg D. Histone methyltransferase activity associated with a human multiprotein complex containing the Enhancer of Zeste protein. Genes Dev. 2002;16:2893–905.

12. Müller J, Hart CM, Francis NJ, Vargas ML, Sengupta A, Wild B, et al. Histone Methyltransferase Activity of a Drosophila Polycomb Group Repressor Complex. Cell. 2002;111:197–208.

13. Gao Z, Zhang J, Bonasio R, Strino F, Sawai A, Parisi F, et al. PCGF Homologs, CBX Proteins, and RYBP Define Functionally Distinct PRC1 Family Complexes. Mol Cell. 2012;45:344–56.

14. Hauri S, Comoglio F, Seimiya M, Gerstung M, Glatter T, Hansen K, et al. A High-Density Map for Navigating the Human Polycomb Complexome. Cell Rep. 2016;17:583–95.

15. Wang H, Wang L, Erdjument-Bromage H, Vidal M, Tempst P, Jones RS, et al. Role of histone H2A ubiquitination in Polycomb silencing. Nature. Nature Publishing Group; 2004;431:873–8.

16. Shao Z, Raible F, Mollaaghababa R, Guyon JR, Wu C, Bender W, et al. Stabilization of Chromatin Structure by PRC1, a Polycomb Complex. Cell. 1999;98:37–46.

17. Francis NJ, Saurin AJ, Shao Z, Kingston RE. Reconstitution of a Functional Core Polycomb Repressive Complex. Mol Cell. 2001;8:545–56.

18. Lo SM, Ahuja NK, Francis NJ. Polycomb Group Protein Suppressor 2 of Zeste Is a Functional Homolog of Posterior Sex Combs. Mol Cell Biol. American Society for Microbiology Journals; 2009;29:515–25.

19. Xu L, Shen W-H. Polycomb Silencing of KNOX Genes Confines Shoot Stem Cell Niches in Arabidopsis. Curr Biol. 2008;18:1966–71.

20. Bratzel F, Yang C, Angelova A, López-Torrejón G, Koch M, Pozo JC del, et al. Regulation of the New Arabidopsis Imprinted Gene AtBMI1C Requires the Interplay of Different Epigenetic Mechanisms. Mol Plant. Elsevier; 2012;5:260–9.

21. Yang C, Bratzel F, Hohmann N, Koch M, Turck F, Calonje M. VAL- and AtBMI1-Mediated H2Aub Initiate the Switch from Embryonic to Postgerminative Growth in Arabidopsis. Curr Biol. 2013;23:1324–9.

22. Morey L, Aloia L, Cozzuto L, Benitah SA, Di Croce L. RYBP and Cbx7 Define Specific Biological Functions of Polycomb Complexes in Mouse Embryonic Stem Cells. Cell Rep. 2013;3:60–9.

23. Tavares L, Dimitrova E, Oxley D, Webster J, Poot R, Demmers J, et al. RYBP-PRC1 Complexes Mediate H2A Ubiquitylation at Polycomb Target Sites Independently of PRC2 and H3K27me3. Cell. 2012;148:664–78.

24. Blackledge NP, Farcas AM, Kondo T, King HW, McGouran JF, Hanssen LLP, et al. Variant PRC1 Complex-Dependent H2A Ubiquitylation Drives PRC2 Recruitment and Polycomb Domain Formation. Cell. 2014;157:1445–59.

25. Endoh M, Endo TA, Endoh T, Isono K, Sharif J, Ohara O, et al. Histone H2A Mono-Ubiquitination Is a Crucial Step to Mediate PRC1-Dependent Repression of Developmental Genes to Maintain ES Cell Identity. PLOS Genet. Public Library of Science; 2012;8:e1002774.

26. Cooper S, Dienstbier M, Hassan R, Schermelleh L, Sharif J, Blackledge NP, et al. Targeting Polycomb to Pericentric Heterochromatin in Embryonic Stem Cells Reveals a Role for H2AK119u1 in PRC2 Recruitment. Cell Rep. 2014;7:1456–70.

27. Illingworth RS, Moffat M, Mann AR, Read D, Hunter CJ, Pradeepa MM, et al. The E3 ubiquitin ligase activity of RING1B is not essential for early mouse development. Genes Dev. 2015;29:1897–902.

28. Blackledge NP, Fursova NA, Kelley JR, Huseyin MK, Feldmann A, Klose RJ. PRC1 Catalytic Activity Is Central to Polycomb System Function. Mol Cell. 2020;77:857–874.e9.

29. Tamburri S, Lavarone E, Fernández-Pérez D, Conway E, Zanotti M, Manganaro D, et al. Histone H2AK119 Mono-Ubiquitination Is Essential for Polycomb-Mediated Transcriptional Repression. Mol Cell. 2020;77:840–856.e5.

30. Pengelly AR, Kalb R, Finkl K, Müller J. Transcriptional repression by PRC1 in the absence of H2A monoubiquitylation. Genes Dev. Cold Spring Harbor Laboratory Press; 2015;29:1487.

31. Kahn TG, Dorafshan E, Schultheis D, Zare A, Stenberg P, Reim I, et al. Interdependence of PRC1 and PRC2 for recruitment to Polycomb Response Elements. Nucleic Acids Res. Oxford Academic; 2016;44:10132–49.

32. Tsuboi M, Kishi Y, Yokozeki W, Koseki H, Hirabayashi Y, Gotoh Y. Ubiquitination-lndependent Repression of PRC1 Targets during Neuronal Fate Restriction in the Developing Mouse Neocortex. Dev Cell. 2018;47:758–772.e5.

33. Zhou Y, Romero-Campero FJ, Gómez-Zambrano Á, Turck F, Calonje M. H2A monoubiquitination in Arabidopsis thaliana is generally independent of LHP1 and PRC2 activity. Genome Biol. 2017;18:69.

34. Kralemann LEM, Liu S, Trejo-Arellano MS, Muñoz-Viana R, Köhler C, Hennig L. Removal of H2Aub1 by ubiquitin-specific proteases 12 and 13 is required for stable Polycomb-mediated gene repression in Arabidopsis. Genome Biol. 2020;21.

35. Yin X, Romero-Campero FJ, de Los Reyes P, Yan P, Yang J, Tian G, et al. H2AK121ub in Arabidopsis associates with a less accessible chromatin state at transcriptional regulation hotspots. Nat Commun. Nature Publishing Group; 2021;12:315.

36. Derkacheva M, Liu S, Figueiredo DD, Gentry M, Mozgova I, Nanni P, et al. H2A deubiquitinases UBP12/13 are part of the Arabidopsis polycomb group protein system. Nat Plants. Nature Publishing Group; 2016;2:1–10.

37. Bowman JL, Kohchi T, Yamato KT, Jenkins J, Shu S, Ishizaki K, et al. Insights into Land Plant Evolution Garnered from the Marchantia polymorpha Genome. Cell. 2017;171:287–304.e15.

38. Kawashima T, Lorković ZJ, Nishihama R, Ishizaki K, Axelsson E, Yelagandula R, et al. Diversification of histone H2A variants during plant evolution. Trends Plant Sci. 2015;20:419–25.

39. Montgomery SA, Tanizawa Y, Galik B, Wang N, Ito T, Mochizuki T, et al. Chromatin Organization in Early Land Plants Reveals an Ancestral Association between H3K27me3, Transposons, and Constitutive Heterochromatin. Curr Biol. 2020;30:573–588.e7.

40. Sanchez-Pulido L, Devos D, Sung ZR, Calonje M. RAWUL: A new ubiquitin-like domain in PRC1 Ring finger proteins that unveils putative plant and worm PRC1 orthologs. BMC Genomics. 2008;9:308.

41. Junco SE, Wang R, Gaipa JC, Taylor AB, Schirf V, Gearhart MD, et al. Structure of the Polycomb Group Protein PCGF1 in Complex with BCOR Reveals Basis for Binding Selectivity of PCGF Homologs. Structure. 2013;21:665–71.

42. Gray F, Cho HJ, Shukla S, He S, Harris A, Boytsov B, et al. BMI1 regulates PRC1 architecture and activity through hom o- and hetero-oligomerization. Nat Commun. Nature Publishing Group; 2016;7:13343.

43. Wong SJ, Gearhart MD, Taylor AB, Nanyes DR, Ha DJ, Robinson AK, et al. KDM2B Recruitment of the Polycomb Group Complex, PRC1.1, Requires Cooperation between PCGF1 and BCORL1. Structure. 2016;24:1795–801.

44. Chittock EC, Latwiel S, Miller TCR, Müller CW. Molecular architecture of polycomb repressive complexes. Biochem Soc Trans. Portland Press; 2017;45:193–205.

45. Lee H-G, Kahn TG, Simcox A, Schwartz YB, Pirrotta V. Genome-wide activities of Polycomb complexes control pervasive transcription. Genome Res. 2015;25:1170–81.

46. Gómez-Zambrano Á, Merini W, Calonje M. The repressive role of Arabidopsis H2A.Z in transcriptional regulation depends on AtBMI1 activity. Nat Commun. Nature Publishing Group; 2019;10:2828.

47. Frangini A, Sjöberg M, Roman-Trufero M, Dharmalingam G, Haberle V, Bartke T, et al. The Aurora B Kinase and the Polycomb Protein Ring1B Combine to Regulate Active Promoters in Quiescent Lymphocytes. Mol Cell. 2013;51:647–61.

48. van den Boom V, Maat H, Geugien M, Rodríguez López A, Sotoca AM, Jaques J, et al. Non-canonical PRC1.1 Targets Active Genes Independent of H3K27me3 and Is Essential for Leukemogenesis. Cell Rep. 2016;14:332–46.

49. Kloet SL, Makowski MM, Baymaz HI, van Voorthuijsen L, Karemaker ID, Santanach A, et al. The dynamic interactome and genomic targets of Polycomb complexes during stem-cell differentiation. Nat Struct Mol Biol. Nature Publishing Group; 2016;23:682–90.

50. Cohen I, Zhao D, Bar C, Valdes VJ, Dauber-Decker KL, Nguyen MB, et al. PRC1 Fine-tunes Gene Repression and Activation to Safeguard Skin Development and Stem Cell Specification. Cell Stem Cell. 2018;22:726–739.e7.

51. Gao Z, Lee P, Stafford JM, von Schimmelmann M, Schaefer A, Reinberg D. An AUTS2–Polycomb complex activates gene expression in the CNS. Nature. Nature Publishing Group; 2014;516:349–54.

52. Linde A-M, Eklund DM, Kubota A, Pederson ERA, Holm K, Gyllenstrand N, et al. Early evolution of the land plant circadian clock. New Phytol. 2017;216:576–90.

53. Ishizaki K, Nishihama R, Ueda M, Inoue K, Ishida S, Nishimura Y, et al. Development of Gateway Binary Vector Series with Four Different Selection Markers for the Liverwort Marchantia polymorpha. PLOS ONE. Public Library of Science; 2015;10:e0138876.

54. Sugano SS, Nishihama R, Shirakawa M, Takagi J, Matsuda Y, Ishida S, et al. Efficient CRISPR/Cas9-based genome editing and its application to conditional genetic analysis in Marchantia polymorpha. PLOS ONE. Public Library of Science; 2018;13:e0205117.

55. Eklund DM, Kanei M, Flores-Sandoval E, Ishizaki K, Nishihama R, Kohchi T, et al. An Evolutionarily Conserved Abscisic Acid Signaling Pathway Regulates Dormancy in the Liverwort Marchantia polymorpha. Curr Biol. 2018;28:3691–3699.e3.

56. Dobin A, Davis CA, Schlesinger F, Drenkow J, Zaleski C, Jha S, et al. STAR: ultrafast universal RNA-seq aligner. Bioinformatics. Oxford Academic; 2013;29:15–21.

57. Love MI, Huber W, Anders S. Moderated estimation of fold change and dispersion for RNA-seq data with DESeq2. Genome Biol. 2014;15:550.

58. Villar CBR, Köhler C. Plant Chromatin Immunoprecipitation. In: Hennig L, Köhler C, editors. Plant Dev Biol Methods Protoc. Totowa, NJ: Humana Press; 2010. p. 401–11. Available from: https://doi.org/10.1007/978-1-60761-765-5_27

59. Bolger AM, Lohse M, Usadel B. Trimmomatic: a flexible trimmer for Illumina sequence data. Bioinformatics. Oxford Academic; 2014;30:2114–20.

60. Langmead B, Salzberg SL. Fast gapped-read alignment with Bowtie 2. Nat Methods. Nature Publishing Group; 2012;9:357–9.

61. Heinz S, Benner C, Spann N, Bertolino E, Lin YC, Laslo P, et al. Simple Combinations of Lineage-Determining Transcription Factors Prime cis-Regulatory Elements Required for Macrophage and B Cell Identities. Mol Cell. 2010;38:576–89.

62. Thorvaldsdóttir H, Robinson JT, Mesirov JP. Integrative Genomics Viewer (IGV): high-performance genomics data visualization and exploration. Brief Bioinform. Oxford Academic; 2013;14:178–92.

63. Ramírez F, Dündar F, Diehl S, Grüning BA, Manke T. deepTools: a flexible platform for exploring deep-sequencing data. Nucleic Acids Res. Oxford Academic; 2014;42:W187–91.

64. Van Bel M, Diels T, Vancaester E, Kreft L, Botzki A, Van de Peer Y, et al. PLAZA 4.0: an integrative resource for functional, evolutionary and comparative plant genomics. Nucleic Acids Res. Oxford Academic; 2018;46:D1190–6.

