## Supplemental information for "H2A ubiquitination is essential for Polycomb Repressive Complex 1-mediated gene regulation in *Marchantia polymorpha*"

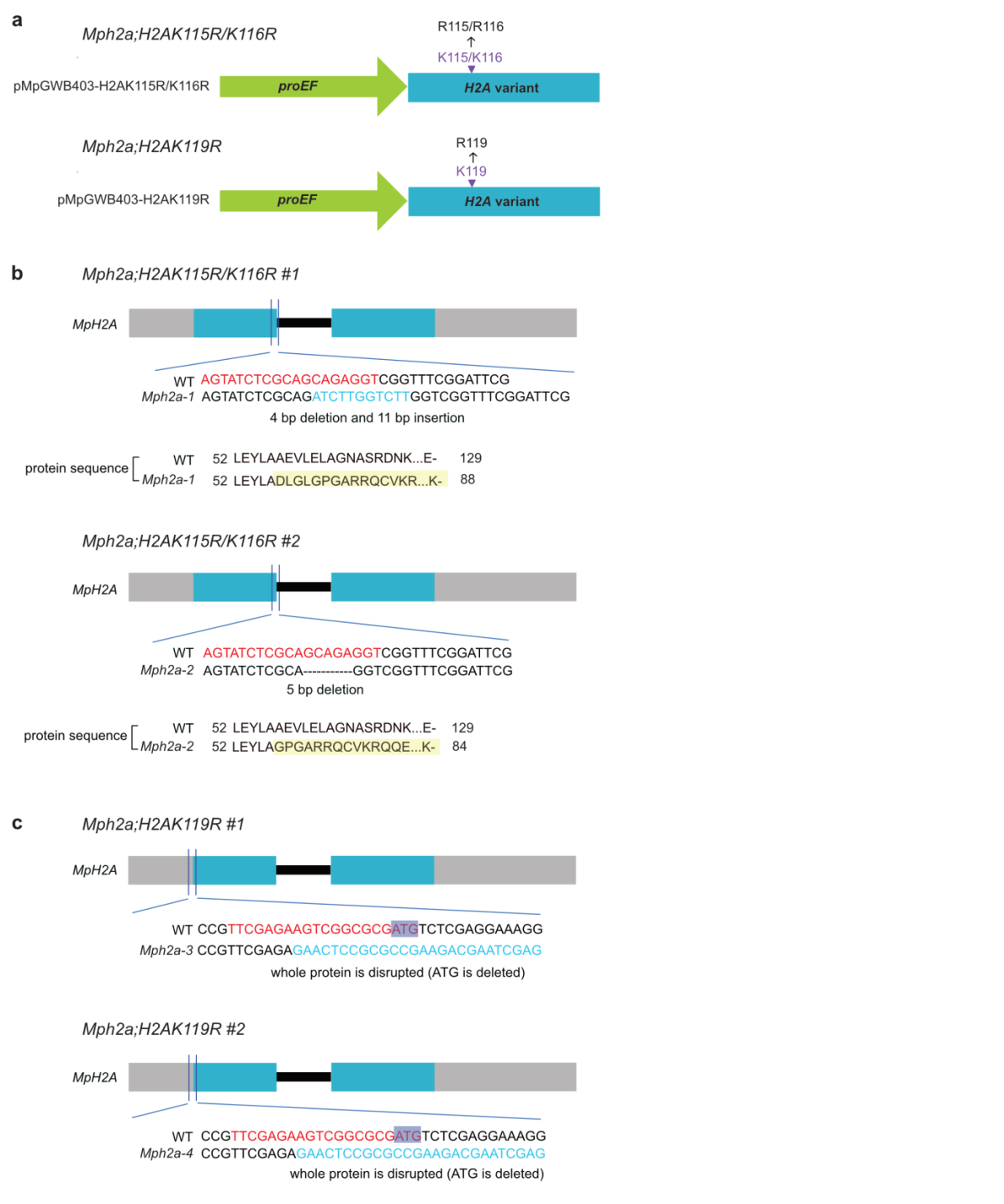

**Fig. S1 Strategy of generating H2Aub depleted lines**

*a.H2A* variants driven by *MpEF* promoter (pMpGWB403-H2AK115R/K116R and pMpGWB403-H2AK119R) and CRISPR/Cas9 constructs with target gRNAs (panel b, c) were co-transformed into *Marchantia* to generate *Mph2a;H2AK115R/K116R* and *Mph2a;H2AK119R* plants. b, c. Disruption of the endogenous *MpH2A* by CRISPR/Cas9. Nucleotide and protein sequence alignments of wild-type (WT) and mutant alleles of MpH2A. *Mph2a;H2AK115R/K116R* #1 refers to *Mph2a-1* expressing *H2AK115R/K116R*, *Mph2a;H2AK115R/K116R* #2 refers to *Mph2a-2* expressing *H2AK115R/K116R*, *Mph2a;H2AK119R* #1 refers to *Mph2a-3* expressing *H2AK119R,* and *Mph2a;H2AK119R* #2 refers to *Mph2a-4* expressing *H2AK119R*. gRNA sequences are shown in red. Newly inserted nucleotides are shown in light blue. Nonsense protein sequences are shaded in yellow. “-” refers to the stop codon. ATG in purple shadow refers to the start codon.

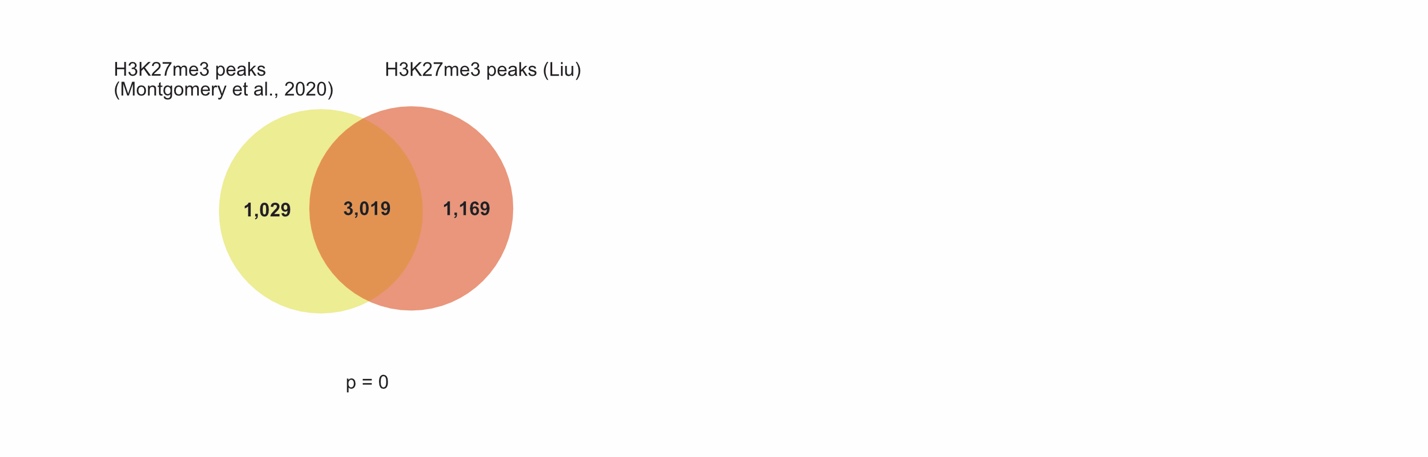

**Fig. S2 Comparison of H3K27me3 data of this study with previously published data.**

Venn diagram showing overlap of wild-type H3K27me3 peaks generated in this study with previously published H3K27me3 peaks (Montgomery et al. 2020). Significance was tested using a Hypergeometric test.

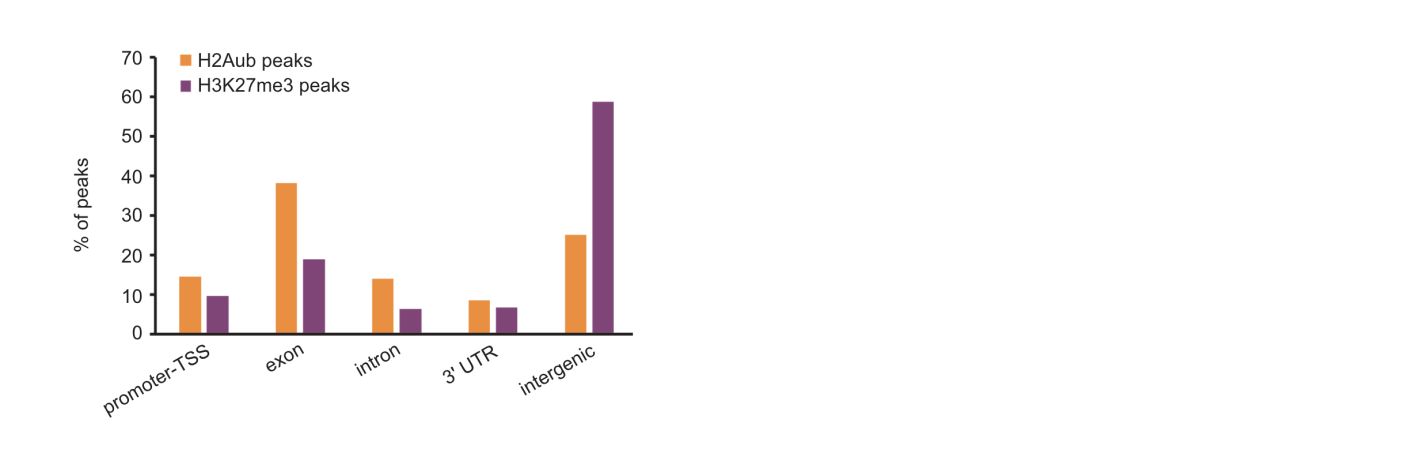

**Fig. S3 Distribution of the H2Aub and H3K27me3 peaks among functional features of the *Marchantia* genome.**

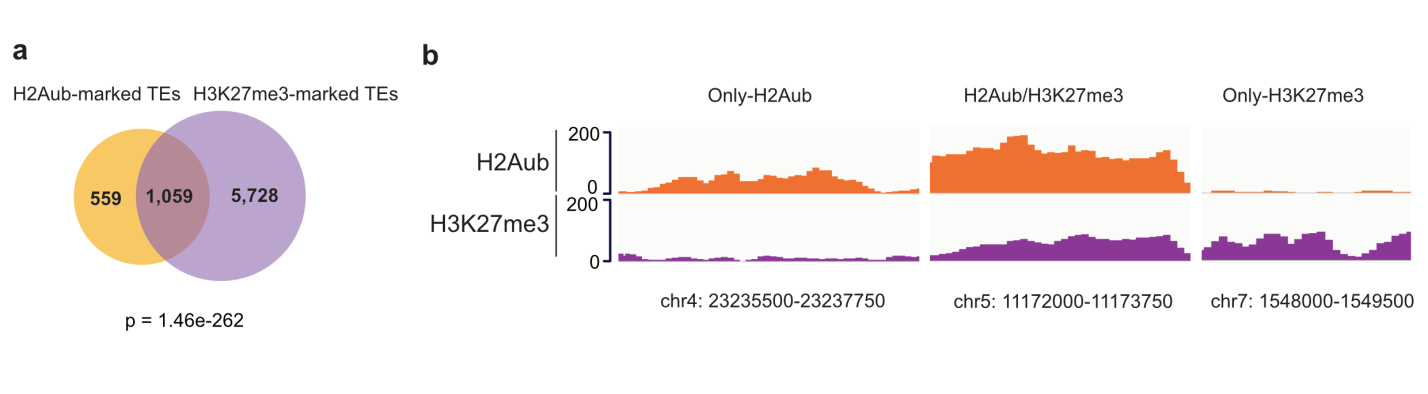

**Fig. S4 The presence of H2Aub and H3K27me3 on transposable elements**

a.Venn diagram showing overlap of H2Aub-marked TEs and H3K27me3-marked TEs. Significance of overlap was tested using a hypergeometric test. b. Genome browser views of selected loci covered by only-H2Aub, H2Aub/H3K27me3 and only-H3K27me3 marks.

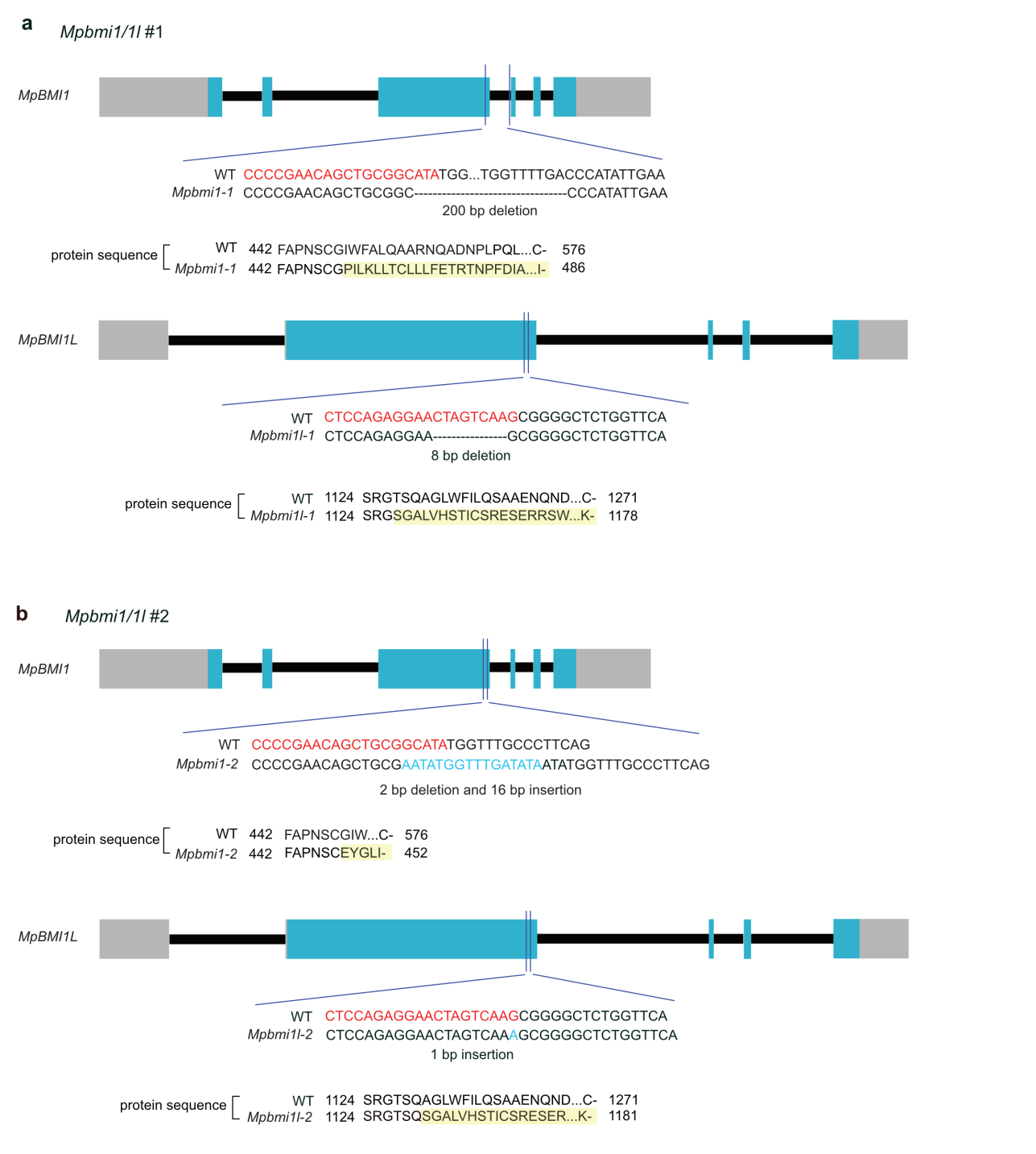

**Fig. S5 Characterization of strong double mutant allele combinations in *MpBMI1/1L***

a, b. Nucleotide and protein sequence alignments of wild-type (WT) and mutant alleles of MpBMI1 and MpBMI1L. *Mpbmi1/1l* #1 (a) contains mutant alleles *Mpbmi1-1* and *Mpbmi1l-1. Mpbmi1/1l* #2 (b) contains mutant alleles *Mpbmi1-2* and *Mpbmi1l-2.* gRNA sequences are shown in red. Newly inserted nucleotides are shown in light blue. Nonsense protein sequences are shaded in yellow. “-” refers to the stop codon.

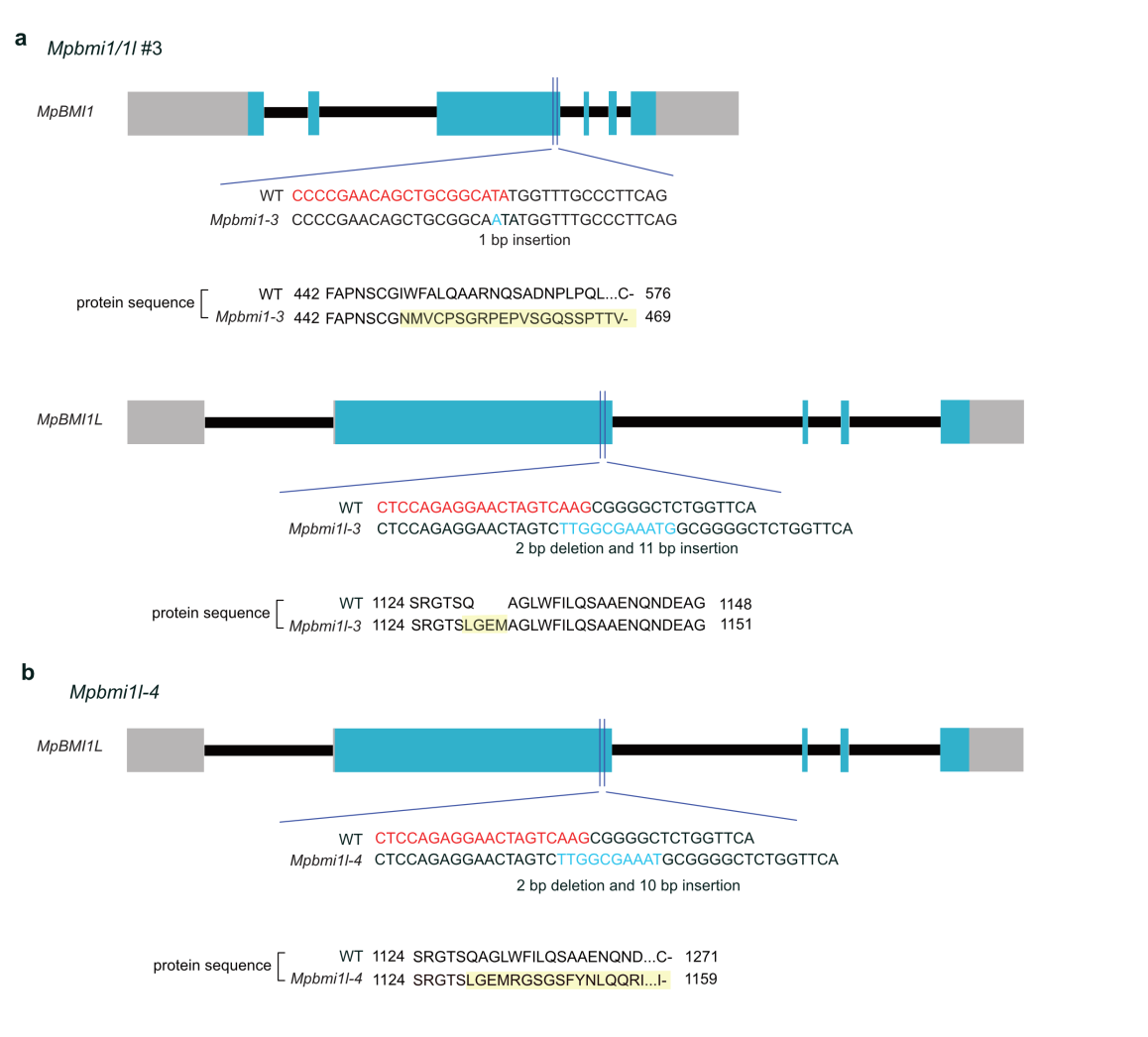

**Fig. S6 Characterization of weak double mutant allele combinations in *MpBMI1/1L***

a, b. Nucleotide and protein sequence alignments of wild-type (WT) and mutant alleles of MpBMI1 and MpBMI1L. *Mpbmi1/1l* #3 (a) contains mutant alleles *Mpbmi1-3* and *Mpbmi1l-3*. *Mpbmi1l-4* (b) is a single mutant. gRNA sequences are shown in red. Newly inserted nucleotides are shown in light blue. Nonsense protein sequences are shaded in yellow. “-” refers to the stop codon.

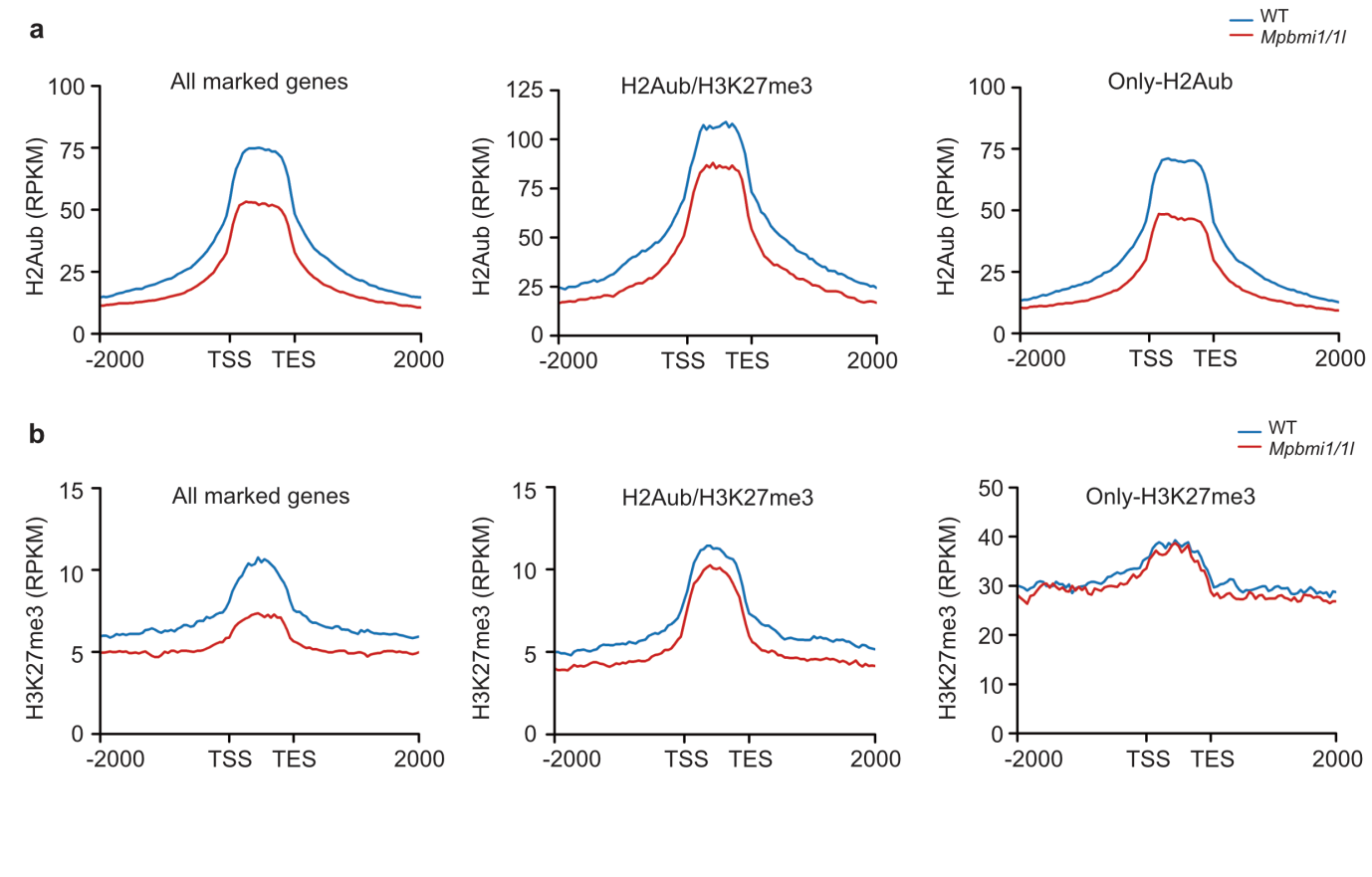

**Fig. S7 H2Aub and H3K27me3 deposition are affected by loss of MpBMI1/1L**

a. Metagene plots showing H2Aub coverage on all marked genes, H2Aub/H3K27me3 genes and only-H2Aub genes in wild type (WT) and *Mpbmi1/1l* mutants. b. Metagene plots showing H3K27me3 coverage on all marked genes, H2Aub/H3K27me3 marked genes and only-H3K27me3 marked genes in WT and *Mpbmi1/1l* mutants.

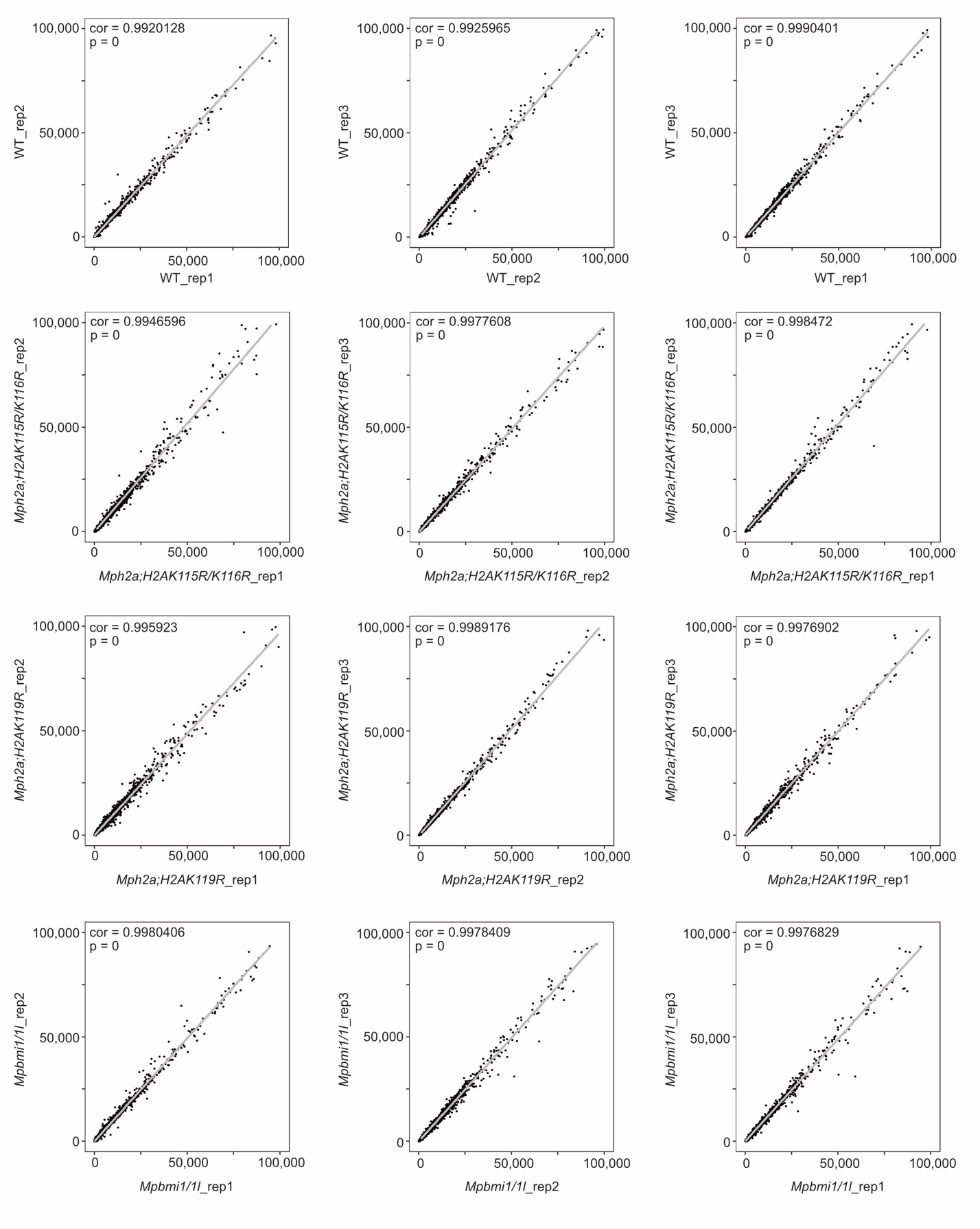

**Fig. S8 Scatter plots comparing RNA-seq triplicates.**

Scatter plots of RPKM (reads per kilobase per million mapped reads) from indicated samples. cor, Pearson correlation coefficient.

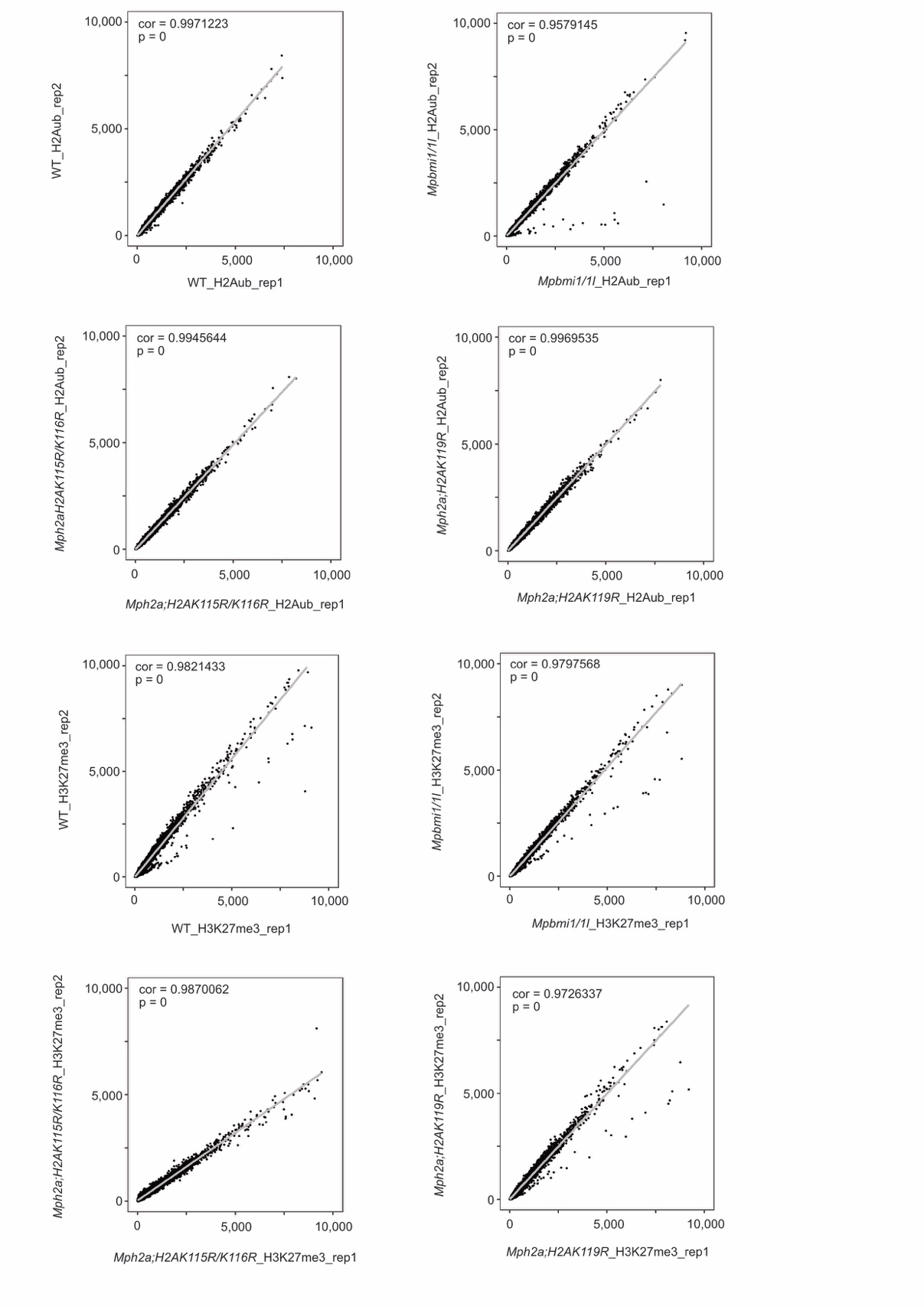

**Fig. S9 Scatter plots comparing ChIP-seq replicates.**

Scatter plots of RPKM (reads per kilobase per million mapped reads) in detected peaks from indicated samples. cor, Pearson correlation coefficient.

| Table S1. Primers used in the constructs. | |
| --- | --- |
| LH4513 | CTCGCCCCGAACAGCTGCGGCATA |
| LH4514 | AAACTATGCCGCAGCTGTTCGGGG |
| LH4517 | CTCGCTCCAGAGGAACTAGTCAAG |
| LH4518 | AAACCTTGACTAGTTCCTCTGGAG |
| LH4012 | CTCGAGTATCTCGCAGCAGAGGT |
| LH4013 | AAACACCTCTGCTGCGAGATACT |
| LH4303 | CTCGCATCGCGCCGACTTCTCGAA |
| LH4304 | AAACTTCGAGAAGTCGGCGCGATG |
| LH3848 | CACCATGTCTCGAGGAAAGGTCAC |
| LH4309 | CTACTCTTTTCCCTCGGCGAGCGAGAGGGCAGGTTTTCCTGACCTCCTGGGCAGCAACACTGAGT |
| LH3849 | CTACTCTTTTCCCTCGGCGAGCGAGAGGGCAGGTCTTCCTGACTTCTTGGGCAG |

| Table S2: Mapping statistics of RNA-seq data | | | | |
| --- | --- | --- | --- | --- |
|  | Biological Replicate | Raw reads | Total aligned | Unique hits |
| WT | R1 | 25621447 | 19858664 | 19400145 |
| WT | R2 | 23965320 | 19421365 | 18952869 |
| WT | R3 | 19021770 | 15144123 | 14797406 |
| *Mph2a;H2AK115R/K116R* | R1 | 21389827 | 16713866 | 16329341 |
| *Mph2a;H2AK115R/K116R* | R2 | 24005264 | 19168652 | 18714591 |
| *Mph2a;H2AK115R/K116R* | R3 | 25236932 | 19682923 | 19222758 |
| *Mph2a;H2AK119R* | R1 | 31767631 | 24714624 | 24135977 |
| *Mph2a;H2AK119R* | R2 | 29951409 | 23012264 | 22493575 |
| *Mph2a;H2AK119R* | R3 | 28751598 | 22140820 | 21616533 |
| *Mpbmi1/1l* | R1 | 30270331 | 23896906 | 23239671 |
| *Mpbmi1/1l* | R2 | 31849730 | 25368771 | 24653253 |
| *Mpbmi1/1l* | R3 | 27844134 | 21864934 | 21271477 |

| Table S3. Mapping statistics of ChIP-seq data | | | | | | |
| --- | --- | --- | --- | --- | --- | --- |
| IP | Genotype | Replicate | Raw reads | Total aligned | Unique hits | coverage (mapped reads*150/haploid genome) |
| H3 | WT | 1 | 21298895 | 15416140 | 8246744 | 16,52 |
| H3 | *Mpbmi1/1l* | 1 | 18258035 | 15638007 | 8207878 | 16,76 |
| H3 | *Mph2a;H2AK115R/K116R* | 1 | 20533143 | 16541500 | 7915436 | 17,72 |
| H3 | *Mph2a;H2AK119R* | 1 | 16720991 | 13652689 | 6790461 | 14,63 |
| H2Aub | WT | 1 | 24791165 | 5751550 | 4211234 | 6,16 |
| H2Aub | *Mpbmi1/1l* | 1 | 25453803 | 4701317 | 3254134 | 5,04 |
| H2Aub | *Mph2a;H2AK115R/K116R* | 1 | 30160712 | 5097069 | 3748722 | 5,46 |
| H2Aub | *Mph2a;H2AK119R* | 1 | 33846077 | 4355990 | 3177103 | 4,67 |
| H3K27me3 | WT | 1 | 65503823 | 7912862 | 2837618 | 8,48 |
| H3K27me3 | *Mpbmi1/1l* | 1 | 63353608 | 7722805 | 2759069 | 8,27 |
| H3K27me3 | *Mph2a;H2AK115R/K116R* | 1 | 76673101 | 6984920 | 2249939 | 7,48 |
| H3K27me3 | *Mph2a;H2AK119R* | 1 | 62512293 | 7545234 | 2633179 | 8,08 |
| H3 | WT | 2 | 20001767 | 15801396 | 8180624 | 16,93 |
| H3 | *Mpbmi1/1l* | 2 | 14887412 | 13102411 | 7068679 | 14,04 |
| H3 | *Mph2a;H2AK115R/K116R* | 2 | 19305042 | 13998086 | 6699146 | 15,00 |
| H3 | *Mph2a;H2AK119R* | 2 | 23105292 | 17465290 | 8623838 | 18,71 |
| H2Aub | WT | 2 | 29465130 | 5333189 | 3895799 | 5,71 |
| H2Aub | *Mpbmi1/1l* | 2 | 28301567 | 5578239 | 3761394 | 5,98 |
| H2Aub | *Mph2a;H2AK115R/K116R* | 2 | 30636755 | 4053243 | 2874614 | 4,34 |
| H2Aub | *Mph2a;H2AK119R* | 2 | 29953615 | 4487052 | 3257408 | 4,81 |
| H3K27me3 | WT | 2 | 54419810 | 7553470 | 2541188 | 8,09 |
| H3K27me3 | *Mpbmi1/1l* | 2 | 48977096 | 12817306 | 4195494 | 13,73 |
| H3K27me3 | *Mph2a;H2AK115R/K116R* | 2 | 43490481 | 7184627 | 2616214 | 7,70 |
| H3K27me3 | *Mph2a;H2AK119R* | 2 | 62046099 | 5484875 | 1827216 | 5,88 |
